# Kinetochore-microtubule attachments are strengthened by Cnn1 stabilization of Stu2

**DOI:** 10.64898/2026.07.13.738289

**Authors:** Nairita Maitra, Devin T. Edwards, Christian R. Nelson, Changkun Hu, Charles L. Asbury, Sue Biggins

**Affiliations:** Howard Hughes Medical Institute, Division of Basic Sciences, Fred Hutchinson Cancer Center, Seattle, WA, United States of America; Department of Physiology and Biophysics, University of Washington, Seattle, WA, United States of America

## Abstract

Accurate chromosome segregation requires kinetochores to form robust, load-bearing attachments to dynamic spindle microtubules, mediated primarily by the Ndc80 complex. Two receptors, Dsn1 and Cnn1 (CENP-T), recruit multiple copies of Ndc80c to the kinetochore. We previously demonstrated that kinetochore components co-purifying with the yeast Dsn1 protein can maintain persistent load-bearing attachments that track with microtubule tip growth and shortening, but the properties of the Cnn1 receptor have not been explored. Using an optical trapping-based assay, we show here that Cnn1 purifications also sustain dynamic microtubule attachments under load. Mutation of a conserved region within the disordered N-terminal tail of Cnn1 weakened attachment strength *in vitro* and caused a growth defect when Dsn1 function was impaired. The Cnn1 mutation reduced Stu2 kinetochore levels without altering other kinetochore proteins. Restoring Stu2, either by direct addition *in vitro* or by tethering it to Ndc80c *in vivo*, rescued both attachment strength and cellular viability. These findings reveal a biophysical role for Cnn1 in enabling Stu2-dependent stabilization of kinetochore-microtubule attachments.

## INTRODUCTION

The transmission of genetic material during cell division relies on interactions between dynamic microtubules and kinetochores, megadalton protein complexes that assemble on chromosomes. The inner kinetochore components (also known as the <u>c</u>onstitutive <u>c</u>entromere-<u>a</u>ssociated <u>n</u>etwork or CCAN) closely associate with the centromeres of chromosomes, while the outer components bind to microtubules (*1-6*). Multiple outer kinetochore subcomplexes mediate interactions between the kinetochore and microtubule tips (*7-9*). The highly conserved Ndc80 complex is particularly important for microtubule binding because it both directly engages with the microtubule and recruits additional microtubule attachment factors (*8, 10-13*).

Ndc80c is a four-protein complex (Ndc80, Nuf2, Spc24, and Spc25) that assembles into a rod-like structure connected by a four-way junction between the Spc24/Spc25 proteins, which provide anchorage to the inner kinetochore, and the Ndc80/Nuf2 proteins, which project outward to interact with the microtubule (*14-17*). Multiple copies of Ndc80c are recruited to the kinetochore via two distinct receptors: Dsn1 (a subunit of the Mis12c) and Cnn1/CENP-T (a subunit of the Cnn1/Wip1 complex), a CCAN component (Fig 1A)(*18, 19*). A conserved motif at the amino terminus of Cnn1/CENP-T binds to the same Spc24-Spc25 domain of Ndc80c that also binds Dsn1 (*18-22*). It is not clear why cells employ two receptors to recruit Ndc80c. One possibility is that these proteins modulate kinetochore-microtubule attachments at different stages of mitosis since Cnn1 is enriched at yeast kinetochores specifically during anaphase (*20, 23*). Another possibility is that the two Ndc80c receptors present the outer kinetochore subcomplexes to the microtubule differently, thereby influencing the strength of microtubule attachments.

**Fig 1.**
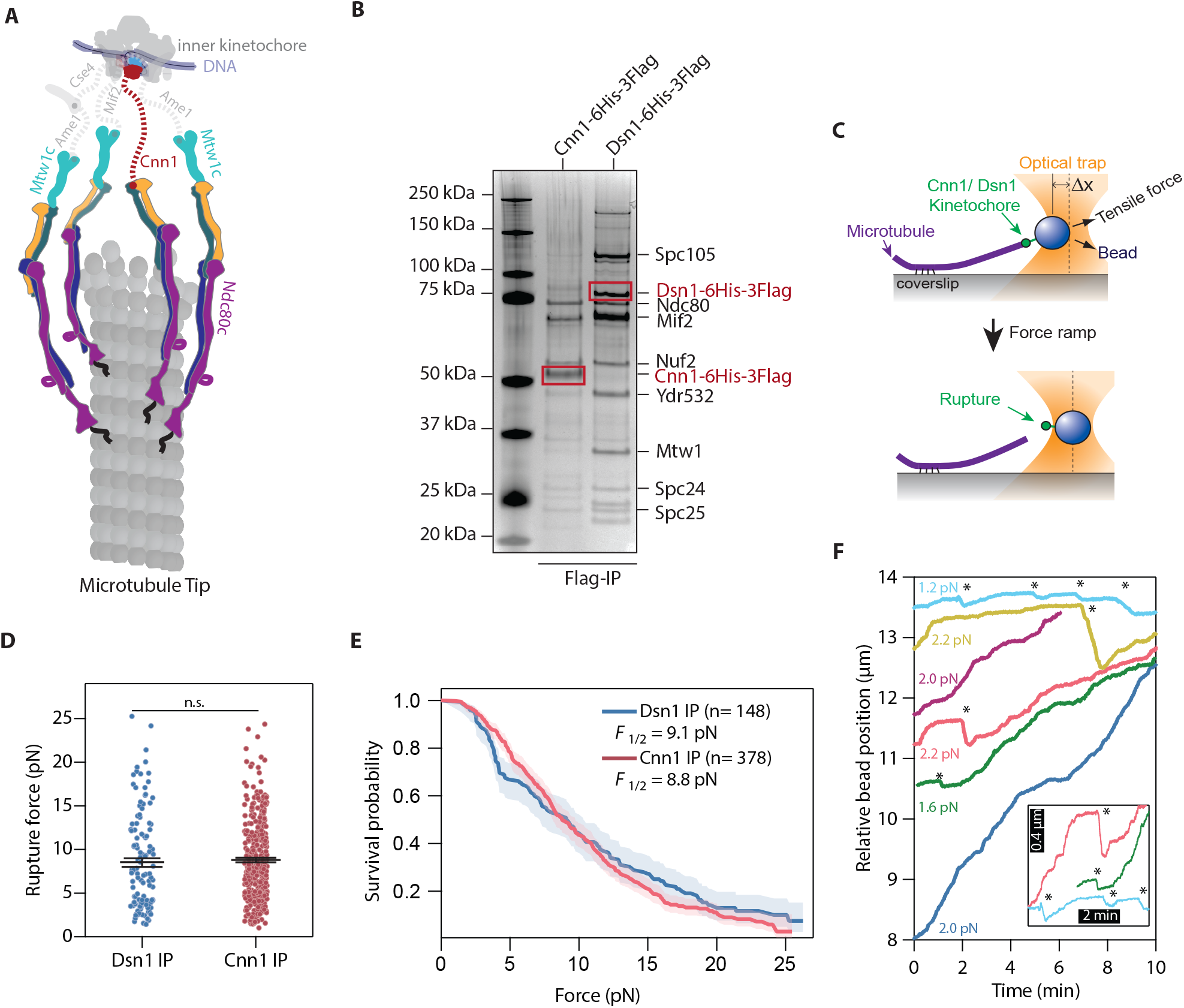
Cnn1 particles maintain microtubule attachments under force. A. Schematic of the kinetochore and microtubule organization in *Saccharomyces cerevisiae*, showing the two Ndc80c receptors, Cnn1 and Mtw1c. B. Cnn1-6His-3Flag (SBY24005) and Dsn1-6His-3Flag (SBY8253) cells were arrested in mitosis with benomyl addition and Cnn1 and Dsn1 were immunoprecipitated and eluted with Flag peptide. The immunoprecipitations were analyzed by silver staining of SDS-PAGE. The red box indicates the immunoprecipitated proteins and the other proteins labeled on the gel were based on prior identifications from (*42*). Molecular weight markers are indicated (kDa, left lane). C. Schematic of the optical trapping assay used to determine the strength of kinetochore-microtubule attachments. Dynamic microtubules were grown from stabilized microtubule seeds attached to a glass coverslip. Purified Cnn1 or Dsn1 particles were linked to polystyrene beads via a 6His tag on Cnn1 or Dsn1. The position of the bead was manipulated with a laser trap and placed at the tip of a growing microtubule until a kinetochore-microtubule attachment was established (top). The force was ramped at 0.25 pN/s until the kinetochore-microtubule interaction was ruptured resulting in the bead detaching from the tip of the microtubule (bottom). D. Scatter plots of rupture force values of Cnn1 and Dsn1 particles. Error bars represent the S.E.M. (n = 123 for Dsn1 and n = 327 for Cnn1 particles where n represents number of rupture events). The p-value =0.16 is calculated from a Mann Whitney test. n.s. represents non-significant. E. The Kaplan-Meier survival probability calculated for bead attachments from Cnn1 (n = 378), and Dsn1 (n = 148) particles shown in (C) where n represents total events (rupture and censored events). Shaded regions represent 95% confidence intervals. P-value = 0.46 calculated from the log-rank test and is non-significant. F. Plot of bead position versus time for Cnn1-purified particle decorated beads held under constant force at the microtubule tip in an optical trap. Traces are offset in space and time for clarity. Positive changes in bead position indicate the bead tracking microtubule growth while decreases in bead position indicate the bead tracking rapid microtubule disassembly (indicated by * in figures). Inset shows an expanded view around 3 transitions from assembly to disassembly.

In addition to engaging the microtubule through its N-terminal globular calponin homology domain and unstructured flexible tail (*11, 16, 24-27*), Ndc80c recruits outer kinetochore complexes that contribute to the load bearing ability of the kinetochore. In yeast, the Dam1 complex (Dam1c; functional ortholog of the human Ska1 complex) and the Stu2 protein (chTOG in human cells) associate with Ndc80c (*28-36*). Dam1c is a heterodecamer that can oligomerize into a ring around the microtubules and associates with the heads of Ndc80/Nuf2 (*35-40*) while Stu2, a member of the conserved XMAP215 family of proteins, directly interacts with the four-way junction of the Ndc80c through its C-terminal segment (CTS) (*30*). However, Stu2 kinetochore localization also depends on its dimerization and TOG1 domains, suggesting that additional features regulate Stu2 kinetochore association through mechanisms that have not been defined (*41*).

Here, we set out to understand the functions of Cnn1 at the kinetochore and to determine whether the biophysical properties of the two Ndc80c receptors differ. Kinetochore particles purified by isolating the Dsn1 receptor from budding yeast have been well characterized and can maintain robust kinetochore-microtubule attachments under forces applied with an optical trap (*42*). However, the biophysical behavior of the Cnn1 linkage to the outer kinetochore has never been explored. To address this, we purified native Cnn1 and found that, like Dsn1, it copurifies outer kinetochore components. The Cnn1-based particles can maintain kinetochore-microtubule attachments that track dynamic microtubule tips under force, similar to Dsn1-based particles. We also discovered that a conserved region of Cnn1 outside of its Ndc80c- and kinetochore-binding domains, named the C-region, contributes to the ability of Cnn1 to mediate kinetochore-microtubule attachments *in vivo* and *in vitro* (*19*). Mutations in the C-region of Cnn1 reduce Stu2 levels at the kinetochore without affecting Ndc80c levels, and cause kinetochore-microtubule attachment defects *in vivo* and *in vitro*. These defects can be rescued by exogenously added Stu2, identifying a previously unrecognized role for Cnn1 in promoting Stu2 localization at the kinetochore.

## RESULTS

### Cnn1-based kinetochore particles form load-bearing attachments to dynamic microtubule tips

Prior work has demonstrated that individual yeast kinetochore particles purified via Dsn1 form load-bearing attachments to dynamic microtubule tips *in vitro* (*42*). The Cnn1 protein is also a receptor for Ndc80, but it is present at low levels in the Dsn1 purifications (fig. S1A, (*43*)). It is unknown whether Cnn1-based purifications can also support microtubule tip-coupling under force or whether their behavior is different from Dsn1 purifications (*18, 19*). To analyze this, we epitope-tagged the endogenous copies of Cnn1 or Dsn1 with a 6His-3Flag tag at their C-termini and purified them from benomyl-arrested mitotic cells by immunoprecipitation (IP) using anti-Flag antibodies. We analyzed the purified material by silver-stained SDS-PAGE (Fig. 1B). As previously reported, Dsn1 copurifies with many kinetochore proteins including the Ndc80 complex (*42*). Ndc80 is also prominent in Cnn1-based purifications, consistent with both Dsn1 and Cnn1 being Ndc80c receptors (Fig. 1B). However, the purifications differed in the number and stoichiometry of other co-purifying proteins, with fewer prominent bands present in the purification based on Cnn1. To further analyze the composition of the purified material, we performed immunoblotting and mass spectrometry. Similar to Dsn1-based purifications, we were able to detect proteins from every kinetochore subcomplex in the Cnn1-purified material (fig. S1A and S1B). However, the proteins that co-purified with Cnn1 were ∼2-3 fold less abundant, making it difficult to assess whether there are differences in the relative protein levels between the two preparations.

To compare the microtubule binding activities of the Cnn1- and Dsn1-based purifications, we utilized an *in vitro* optical trapping assay (*42, 44*). Either the Cnn1 or Dsn1 protein and its copurifying components were linked to polystyrene beads via the -6His tag and introduced into a slide-channel containing dynamic microtubules grown from coverslip-anchored seeds. Individual beads were held in the trap and attached to the tip of a single microtubule to reconstitute kinetochore-microtubule tip-coupling (Fig. 1C, top). As demonstrated previously for Dsn1-based kinetochore particles, the material purified via Cnn1 supported specific binding to dynamic microtubules (fig. S2A) (*42*). We measured the strength of attachment by gradually increasing the force across the bead-microtubule interface (at a fixed loading rate of 0.25 pN/s). The force at which the bead detached from the microtubule was defined as the rupture force (Fig. 1C, bottom). Sampling rupture forces for many kinetochore particles allowed us to assess mean strengths (Fig. 1D), which were 8.5 ± 0.5 pN (mean ± S.E.M.) for Dsn1-based particles and 8.8 ± 0.3 pN (mean ± S.E.M.) for beads decorated with the Cnn1-purified material. For both samples, a small fraction of censored events occurred when the interaction strength exceeded the maximum escape force of the trap, the microtubule detached from the surface, or the ramp was otherwise interrupted before detachment. To include both ruptures and censored events in our analysis, we calculated the Kaplan-Meier survival probability as a function of force (Fig. 1E). We denote the force at which 50% of the kinetochore-microtubule attachments survive as *F*_1/2._ The survival distribution for Dsn1-purified particles was broad, with an *F*_1/2_ of 9.1 pN (Fig. 1E, 95% CI: 7.1 pN – 10.3 pN), consistent with prior measurements (*31, 42, 45*). Cnn1 particles showed a remarkably similar survival distribution with an *F*_1/2_ of 8.8 pN (Fig. 1E, 95% CI: 8.1 pN – 9.7 pN). To test whether these attachment strengths represented the activity of individual particles, we varied the concentration of Cnn1-purified material bound to the beads. Reducing the amount of Cnn1-purified material lowered the fraction of beads that bound microtubules, but the strength of the resulting attachments did not change (fig. S2B), confirming single-particle activity. We verified that the microtubule binding activity of Cnn1-based material required the Ndc80c because particles purified from an *ndc80-1* temperature sensitive mutant showed almost no binding (fig. S2C and S2D). We also performed force-clamp experiments where Cnn1-purified particles were held under constant force at dynamic microtubule tips (Fig. 1F). Beads tracked with the microtubule tip during periods of both microtubule assembly and disassembly, similar to Dsn1-purified particles (*42*). These results show that Cnn1 purified particles carry outer kinetochore components and maintain load-bearing attachments to dynamic microtubule tips, establishing a new platform to further interrogate Cnn1 functions.

### The conserved Cnn1 C-region is important for kinetochore function when Mis12c is crippled

Cnn1 contains a histone fold domain (HFD) at its C-terminus in addition to the Ndc80 binding domain in its N-terminus (Fig. 2A) (*18*). There is also a short hydrophobic segment (157-163) in the disordered N-terminal tail that is highly conserved across *Saccharomyces* species (Fig. 2A, bottom). This patch is called the C-region, and prior work suggests it is important for Cnn1 kinetochore function (*19*). In addition, a recent crosslinking mass spectrometry study shows that amino acids near the C-region might interact with Spc24/Spc25 components of the Ndc80c, suggesting that the C-region could serve as a contact point for outer kinetochore components (*46*).

**Fig 2.**
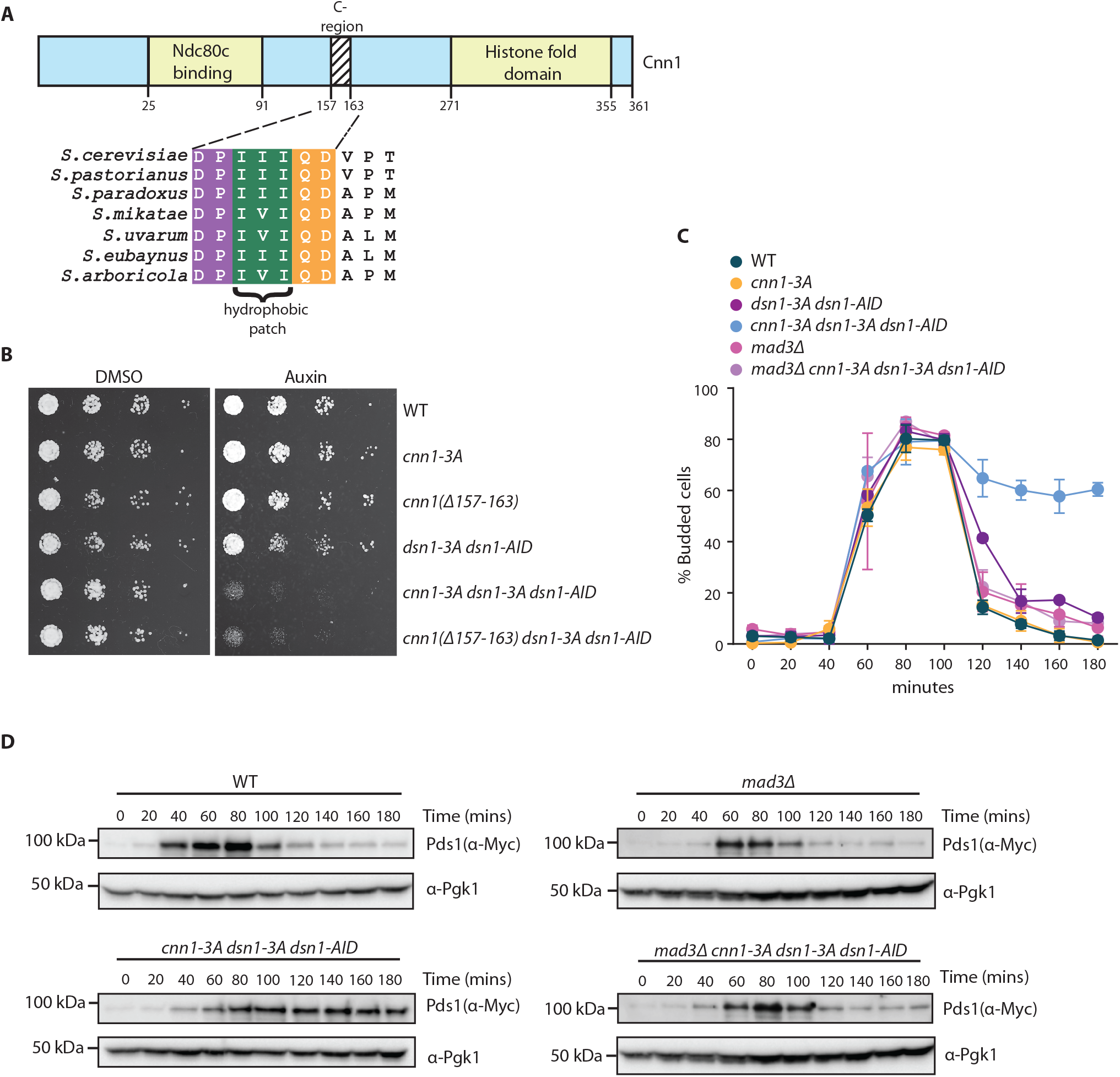
Mutations in the conserved C-region of Cnn1 activate the spindle assembly checkpoint in *dsn1-3A* cells. A. Schematic of Cnn1 domain organization. Alignment of the C-region (amino acids 157 to 163) in different budding yeast species with a hydrophobic patch highlighted (bottom). B. Five-fold serial dilutions of wild type (SBY24005), *cnn1-3A* (SBY24006), *cnn1(Δ157-163)* (SBY24552), *dsn1-3A dsn1-AID* (SBY22977), *cnn1-3A dsn1-3A dsn1-AID* (SBY22972), and *cnn1(Δ157-163) dsn1-3A dsn1-AID* (SBY24652) cells were plated on YPD with DMSO or 500 μM auxin. Cells were incubated at 23 °C and images were taken 48 hours after plating. C. Budding index (% budded cells) was measured for the cells in (A) along with *mad3Δ* (SBY24391), and *mad3Δ cnn1-3A dsn1-3A dsn1-AID* (SBY24389) cells. Cells were arrested in G1 with alpha factor for three hours and auxin was added for the last 30 minutes. The cells were then released into YPD containing auxin (time zero on graph). The percentage of budded cells was monitored every 20 minutes. The mean percentage of budded cells from at least two independent experiments (n > 200 cells per time point) is plotted, with error bars representing the standard deviation. D. Wild type (SBY24387), *cnn1-3A dsn1-3A dsn1-AID* (SBY24549), *mad3Δ* (SBY24738), *mad3Δ cnn1-3A dsn1-3A dsn1-AID* (SBY24789) cells containing Pds1-myc were arrested in G1 with alpha factor for three hours and auxin was added for the last 30 minutes. The cells were then released into YPD containing auxin (time zero). The cells were harvested every 20 minutes after G1 release. Pds1-myc levels were assayed via immunoblot (n=2). Pgk1 levels were used as loading control.

To further explore Cnn1’s role in kinetochore-microtubule attachments, we mutated the C-region hydrophobic patch of three isoleucines to three alanines (Cnn1-3A) or completely deleted the C-region from amino acids 157 to 163 (Cnn1(Δ157-163)). We confirmed that mutation or deletion of the C-region did not alter Cnn1 protein levels or kinetochore localization (fig. S3A-3C). Unlike other species, Cnn1 is non-essential in budding yeast but becomes critical when the Mis12c pathway is crippled by a Dsn1 phosphomutant (*dsn1-3A*) (fig. S4) (*18, 20, 47*). To test whether the C-region is important for kinetochore function *in vivo*, we therefore crossed the *cnn1* mutants to a *dsn1-3A* strain. All *dsn1-3A* strains contained an auxin-inducible degron (*dsn1-AID*) at the endogenous locus to control protein levels and the *dsn1-3A* allele integrated at an ectopic locus. We assessed the viability of wild type, *cnn1-3A*, *cnn1(Δ157-163)*, *dsn1-3A*, *cnn1-3A dsn1-3A,* and *cnn1(Δ157-163) dsn1-3A* cells on media containing either auxin or DMSO (control). Both *cnn1* mutants were synthetically sick with the *dsn1-3A* allele (Fig. 2B), indicating that the C-region is critical for Cnn1 function. Since *cnn1-3A* and *cnn1(Δ157-163)* cells grew similarly in this assay, we proceeded to study just the *cnn1-3A* mutant *in vivo* and *in vitro*.

To determine why the *cnn1-3A dsn1-3A* cells are sick, we analyzed cell cycle progression. Wild type, *cnn1-3A*, *dsn1-3A*, and *cnn1-3A dsn1-3A* cells were released from a G1 arrest into auxin-containing media to degrade Dsn1-AID and monitored for cell cycle progression by budding index. Although the strains budded with similar kinetics, the *cnn1-3A dsn1-3A* cells remained large budded while the other strains exited the cell cycle and became unbudded (Fig. 2C). These data suggest that the *cnn1-3A dsn1-3A* cells are arrested in mitosis due to activation of the spindle assembly checkpoint, the surveillance system that halts the cell cycle when there are defects in kinetochore-microtubule attachments (*48, 49*). To test this, we deleted the *MAD3* checkpoint gene and found that *mad3Δ cnn1-3A dsn1-3A* cells progressed through the cell cycle like wild type, consistent with spindle checkpoint activation in the *cnn1-3A dsn1-3A* cells (Fig. 2C).

To determine whether the spindle assembly checkpoint was triggered, we analyzed the levels of the Pds1 anaphase inhibitor (*50*). Pds1 accumulates until all kinetochores make proper microtubule attachments at metaphase and is then degraded to allow cells to progress to anaphase (*51*). We arrested wild type, *cnn1-3A dsn1-3A*, *mad3Δ,* and *mad3Δ cnn1-3A dsn1-3A* cells containing Pds1-myc in G1 and released them into auxin. As expected, Pds1 levels cycled in wild type and *mad3Δ* cells (Fig. 2D). In contrast, the *cnn1-3A dsn1-3A* cells maintained high levels of Pds1 throughout the time course that were dependent on the *MAD3* spindle assembly checkpoint gene (Fig. 2D). Consistent with this, the *cnn1-3A dsn1-3A* cells were inviable when the *MAD3* gene was deleted (fig. S5). Taken together, these data suggest that there are impaired kinetochore-microtubule attachments in *cnn1-3A dsn1-3A* cells and that the spindle assembly checkpoint provides time for some of them to be corrected.

### The Cnn1 C-region mutants exhibit defective kinetochore clustering in *dsn1-3A* cells

To directly monitor kinetochore-microtubule attachments in the *cnn1-3A dsn1-3A* cells, we fluorescently tagged the Mtw1 kinetochore protein with mCherry and the microtubules with Tub1-GFP. During metaphase, budding yeast kinetochores cluster when they biorient, appearing as two foci (*2, 52-54*) (Fig. 3A). As the cells enter anaphase, the kinetochore foci separate into the mother and daughter cells. To analyze kinetochores, we arrested wild type, *cnn1-3A, dsn1-3A,* and *cnn1-3A dsn1-3A* cells in G1, released them into auxin and analyzed the cells that were in metaphase as defined by short spindles. In wild type and single mutant cells, most metaphase cells exhibited two kinetochore clusters, indicating proper bioriented attachments (Fig. 3B, 3C). In contrast, 24 ± 3% (mean ± S.D.) of the *cnn1-3A dsn1-3A* cells had multiple kinetochore foci that were declustered along the spindle axis, indicating defective kinetochore-microtubule attachments (Fig. 3B, 3C). We note that approximately 6 ± 2% of wild type cells also exhibited defective kinetochore clustering, likely due to the cells containing epitope-tags on both kinetochores and tubulin.

**Fig 3.**
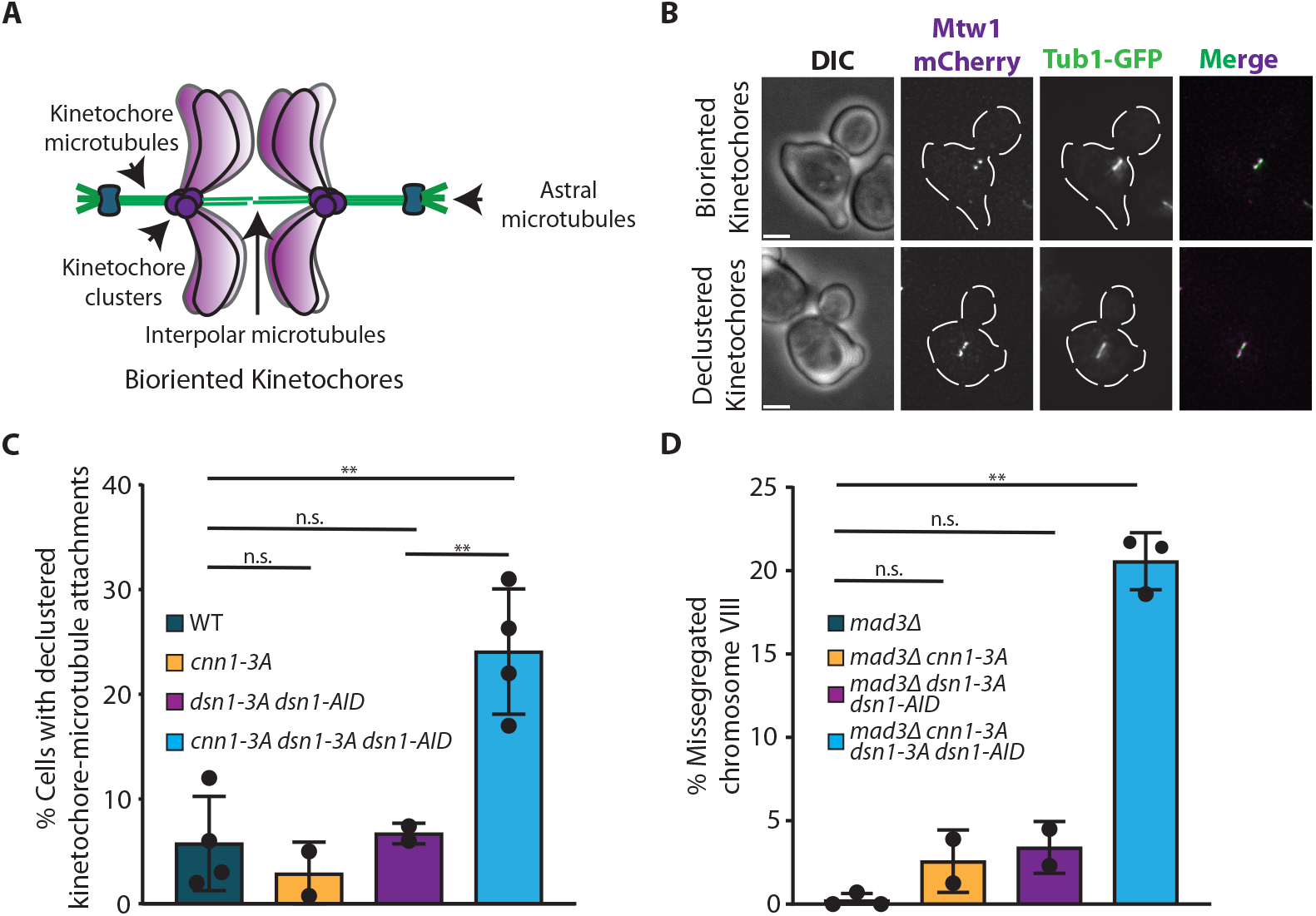
*Cnn1-3A dsn1-3A* mutant cells exhibit kinetochore-microtubule attachment defects. A. Schematic of bioriented kinetochore clusters in budding yeast during metaphase shows they cluster into two foci. B. Representative images of either bioriented clustered kinetochores (top panel) or declustered kinetochores (bottom panel) using Mtw1-mcherry (shown in magenta) as the kinetochore marker. Tub1-GFP, shown in green, was used to identify the spindle. Scale bar is 2 μm. C. Wild type (SBY21592), *cnn1-3A* (SBY21908), *dsn1-3A dsn1-AID* (SBY22830), and *cnn1-3A dsn1-3A dsn1-AID* (SBY22831) cells were arrested in G1 with alpha factor for three hours and auxin was added for the last 30 minutes. The cells were then released into YPD containing auxin. Cells were harvested 75 and 80 minutes after release during metaphase to analyze kinetochores. The percentage of cells with declustered kinetochore-microtubule attachments in each strain during metaphase was quantified across at least 2 experiments (n > 300 total cells per strain). The error bar represents standard deviation. The p-value was determined using Welch’s t-test. The p-value between WT vs *cnn1-3A* is 0.481; WT vs *dsn1-3A dsn1-AID* is 0.7102; WT vs *cnn1-3A dsn1-3A dsn1-AID* is 0.0033; *dsn1-3A dsn1-AID* vs *cnn1-3A dsn1-3A dsn1-AID* is 0.0084. ** represents <0.01. n.s. represent non-significant. D. *Mad3Δ* (SBY24542), *mad3Δ cnn1-3A* (SBY24544), *mad3Δ dsn1-3A dsn1-AID* (SBY24336), and *mad3Δ cnn1-3A dsn1-3A dsn1-AID* (SBY24565) cells were arrested in G1 with alpha factor for three hours and auxin was added for the last 30 minutes. The cells were then released into YPD containing auxin and harvested 100 mins after the release to obtain cells in anaphase. The percentage of cells with missegregated chromosomes is graphed. The error bar represents the standard deviation of at least 2 independent experiments (n > 200 total cells for each strain). The p-value was calculated using the Mann-Whitney test. The p-value calculated between WT vs *cnn1-3A* is 0.2; WT vs *dsn1-3A dsn1-AID* is 0.1386; WT vs *cnn1-3A dsn1-3A dsn1-AID* is 0.0015. ** represents <0.01; n.s. represents non-significant.

We next tested whether the impaired kinetochore-microtubule attachments in the *cnn1-3A dsn1-3A* cells lead to chromosome mis-segregation. We fluorescently marked chromosome VIII by integrating lactose operators near the centromere (at QCR10, 2.1 kb from *CEN8*) in cells expressing a LacI-GFP fusion protein (*55*). We also deleted *MAD3* to ensure cell cycle progression. We released *mad3Δ, mad3Δ cnn1-3A, mad3Δ dsn1-3A,* and *mad3Δ cnn1-3A dsn1-3A* cells from G1 into media containing auxin and monitored chromosome VIII segregation in anaphase cells as indicted by DAPI-labeled DNA masses in the mother and bud. Nearly all *mad3Δ* (1 ± 0.2%), *mad3Δ cnn1-3A* (3 ± 1.3%), and *mad3Δ dsn1-3A* (4 ± 1.1%) (mean percent mis-segregated ± S.D.) cells properly segregated chromosome VIII. However, *mad3Δ cnn1-3A dsn1-3A* cells exhibited significant chromosome mis-segregation with 21 ± 0.98 % of anaphase cells containing GFP signal in only one of the two nuclei (Fig. 3D). Chromosome mis-segregation can be due to defects in error correction or kinetochore-microtubule attachments. These defects can be distinguished by determining whether the chromosomes move to the bud (due to error correction defects that lead to hyper-stable microtubule attachments) or remain in the mother (due to weak microtubule attachments) (*56*). We therefore analyzed where the chromosomes localized to determine the nature of the chromosome missegregation events. The majority of missegregation events resulted in both sister chromatids remaining in the mother cell (90 ± 1.0%), consistent with the *cnn1-3A dsn1-3A* cells having weak kinetochore-microtubule attachments that lead to chromosome missegregation.

### The C-region is important for the attachment strength of Cnn1-based kinetochore particles

We next asked whether the Cnn1 C-region affects kinetochore-microtubule attachments *in vitro*. Cnn1 or Cnn1-3A were purified from benomyl-arrested mitotic cells and the co-purifying material appeared similar in composition as assessed by silver stain gel analysis (Fig. 4A). We coupled the purified material to beads and measured rupture forces in the optical trapping assay. Like wild type Cnn1 particles, the rupture forces of the Cnn1-3A particles did not depend strongly on the concentration of purified material bound to the beads (fig. S6), consistent with single-particle-based interactions. Although the Cnn1-3A particles were able to bind stably to the tips of dynamic microtubules, the interactions ruptured at significantly weaker forces with an *F*_1/2_ of 4.8 pN (95% CI: 4.2 pN –5.2 pN), compared to 8.8 pN for wild type Cnn1 particles (95% CI: 8.1 pN – 9.7 pN) (Fig. 4B). The lower rupture strength of Cnn1-3A particles, only about half that of wild type, is similar to previous measurements of Dsn1-based particles deficient in outer kinetochore components (*31, 42*). The Cnn1 C-region is therefore critical to the attachment strength of the Cnn1-based kinetochore particles, possibly by promoting outer kinetochore assembly.

**Fig 4.**
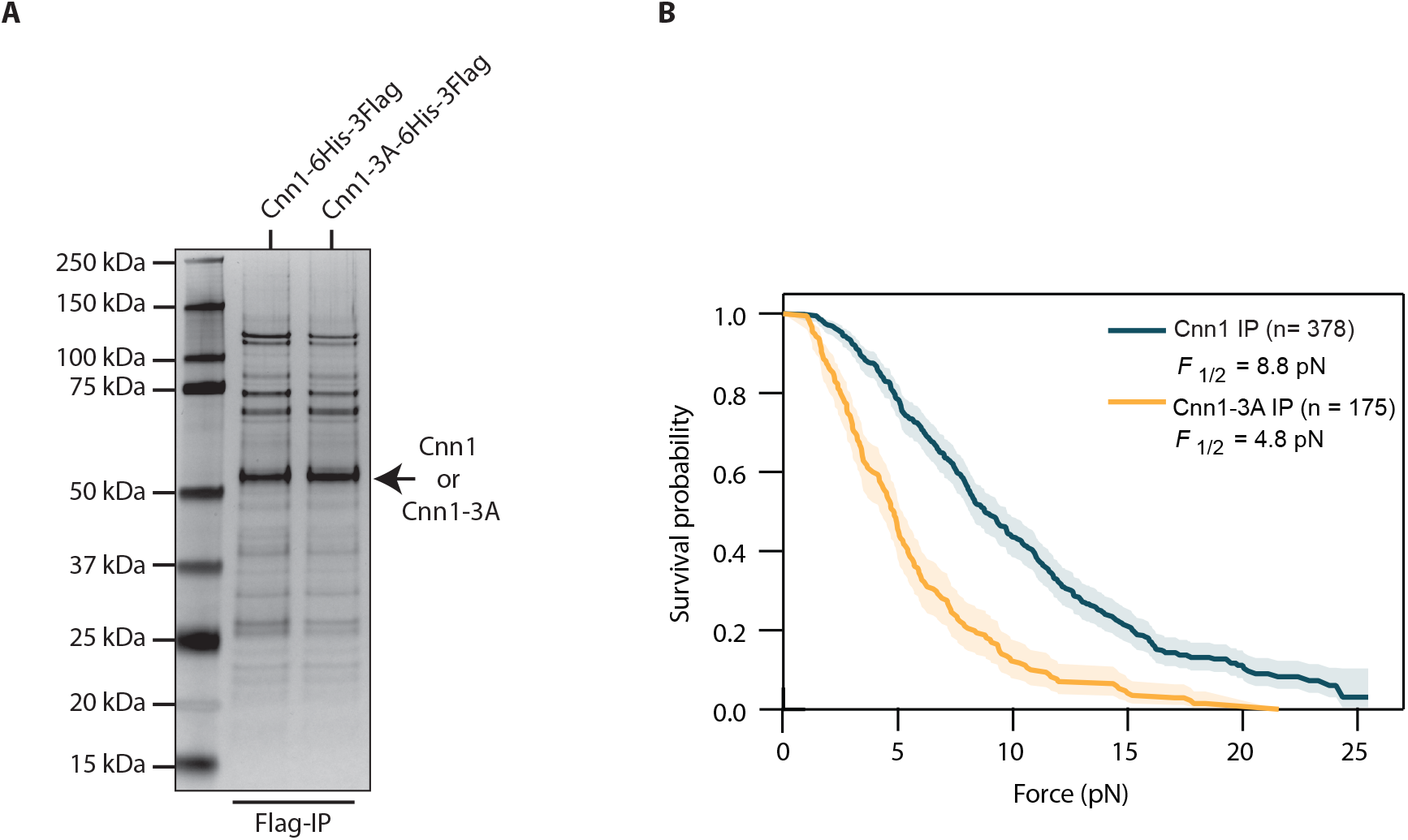
Mutation of the Cnn1 C-region weakens kinetochore-microtubule attachments *in vitro.* A. Purifications of Cnn1-6His-3Flag (SBY24005) and Cnn1-3A-6His-3Flag (SBY24006) cells arrested in mitosis via benomyl treatment were analyzed by silver staining of SDS-PAGE. The arrow indicates the Cnn1 proteins. B. The Kaplan-Meier survival probability of bead attachment for rupture force experiments of Cnn1-6His-3Flag (n = 378) and Cnn1-3A-6His-3Flag (n = 175) particles where n represents the number of total events (rupture and censored events). Shaded regions represent 95% confidence intervals. The p-value calculated from the log-rank test is 2e-21. (Wild type Cnn1 data obtained from the experiments shown in Figure 1C and 1D).

### The Cnn1 C-region promotes Stu2’s association with kinetochores

We further explored the composition of the Cnn1-3A purifications to determine the underlying defects. Although the silver-stain analysis of SDS-PAGE did not exhibit any obvious differences (Fig. 4A), some kinetochore proteins were sub-stoichiometric and not visible, so we performed immunoblotting. Although we did not detect changes in Ndc80 levels, we observed a variable decrease in Stu2 levels in Cnn1-3A purifications that was more apparent when Stu2 was epitope tagged (Stu2-3V5) (Fig. 5A, fig. S7).

**Fig 5.**
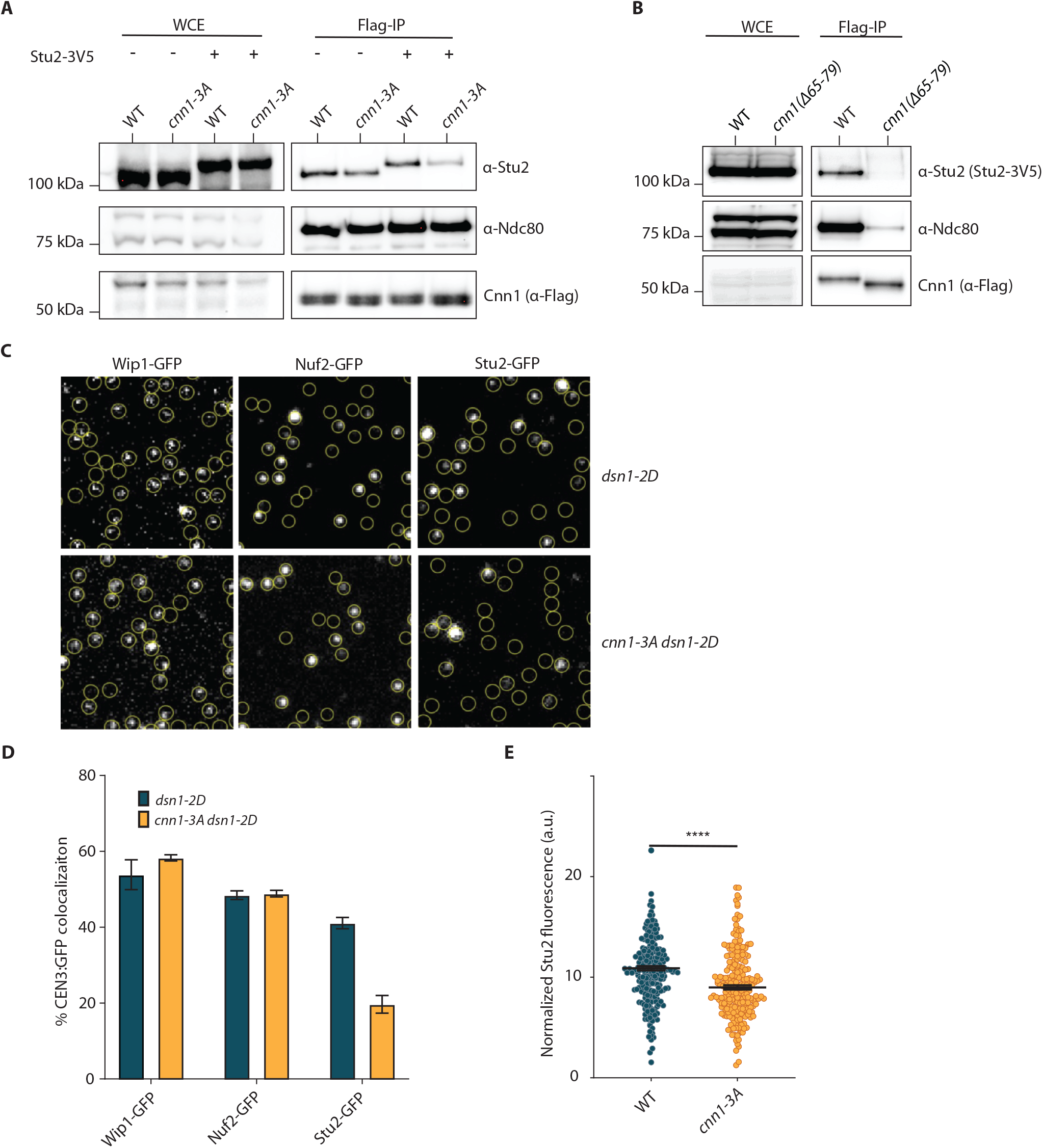
The Cnn1 C-region promotes Stu2 association with the kinetochore. A. Cnn1-3Flag was immunoprecipitated from cells arrested in mitosis via benomyl treatment. Ndc80 and Stu2 levels were analyzed via immunoblot. Strains expressing Stu2 (Cnn1-3Flag: SBY11586 and Cnn1-3A-3Flag: SBY22078) or Stu2-3V5 (Cnn1-3Flag: SBY21682 and Cnn1-3A-3Flag: SBY21805) were used. B. Cnn1-3Flag was immunoprecipitated from benomyl-arrested cells expressing either wildtype Cnn1-6His-3Flag (SBY24923) or Cnn1(Δ65-79)-6His-3Flag (SBY25530) with Stu2-3V5. Ndc80 and Stu2 levels were analyzed via immunoblot. C. Representative TIRFM images of Wip1-GFP, Nuf2-GFP, or Stu2-GFP localized to surface tethered *CEN3* DNA molecules (location shown by yellow circles) after incubation for 180 minutes with lysates from strains expressing *WIP1-GFP dsn1-2D* (SBY22205), *WIP1-GFP dsn1-2D cnn1-3A* (SBY24956), *NUF2-GFP dsn1-2D* (SBY23258), *NUF2-GFP dsn1-2D cnn1-3A* (SBY24955), *STU2-GFP dsn1-2D* (SBY22133) and *STU2-GFP dsn1-2D cnn1-3A* (SBY24954). Scale bar is 5 μm. D. Percentage of Atto-647 tagged *CEN3* DNA colocalized with Wip1-GFP or Nuf2-GFP or Stu2-GFP after 180 minutes of incubation with lysates from above mentioned cells, as measured by single molecule TIRF assays. 3000 DNA molecules were imaged at each time point for each strain and biological replicate (n = 3). The p-value calculated from t-test over three experiments at 180 mins. The p-value for *WIP1-GFP dsn1-2D* vs *WIP1-GFP dsn1-2D cnn1-3A* is 0.044; *NUF2-GFP dsn1-2D* vs *NUF2-GFP dsn1-2D cnn1-3A* is 0.8715; *STU2-GFP dsn1-2D* vs *STU2-GFP dsn1-2D cnn1-3A* is <0.0001. E. Quantification of Stu2 levels at the kinetochore *in vivo*. Wild type (SBY24782) and *cnn1-3A* (SBY24741) cells containing Stu2-GFP were released from G1 and harvested after 75 mins to obtain metaphase cells. The fluorescence intensity of Stu2-GFP was quantified and normalized to the Ame1-mKate intensity a.u. is arbitrary units.. The p-value is calculated from the Mann-Whitney test across three experiments (n > 200 total cells) and is <0.01.

Because Ndc80c is the known Stu2 kinetochore receptor, we were surprised that the C-region of Cnn1 influenced Stu2 kinetochore levels (*30, 31*). We confirmed that Stu2 association with Cnn1 purifications is through Ndc80 by immunoprecipitating a Cnn1 mutant protein that cannot interact with Ndc80. We purified Cnn1 and the Cnn1(Δ65-79) mutant that eliminates the Ndc80 interaction site and analyzed the levels of co-purifying Ndc80 and Stu2 (Fig. 5B). As expected, Ndc80 association was abolished in the Cnn1(Δ65-79) purification. All detectable Stu2 was also eliminated, confirming that Stu2 association with Cnn1 requires Ndc80c.

To quantitatively assess Ndc80 and Stu2 levels in the *cnn1-3A* mutant, we turned to a total internal <u>r</u>eflection <u>f</u>luorescence (TIRF)-based kinetochore assembly assay that monitors assembly kinetics for kinetochore proteins on single centromeric DNA molecules (*57, 58*). In this assay, outer kinetochore assembly is not robust (*58*). To enhance it, we used lysates from cells carrying phospho-mimetic aspartic acid substitutions at Dsn1 S240 and S250 (*dsn1-2D*) in place of *dsn1-3A*. We confirmed that in the Dsn1-2D background, Cnn1-3A still co-purified less Stu2 than wild type Cnn1 particles (fig. S8A). To perform the assay, Atto-647-labeled DNA molecules containing the centromere of chromosome III (*CEN3* DNAs) were sparsely attached to glass coverslips, enabling direct observation of the individual *CEN3* molecules. Next, yeast lysates from benomyl arrested cells carrying either fluorescent-tagged Cnn1 (visualized via Wip1-GFP, the obligate Cnn1 partner (*58*)), Nuf2-GFP, or Stu2-GFP were flowed onto the coverslips. After incubating for different durations (0, 30, 60, 90, 120, 150 or 180 minutes), the lysate was washed away before imaging. We then quantified the fraction of CEN3 DNAs that were co-localized with Wip1-GFP, Nuf2-GFP, or Stu2-GFP (Fig. 5C-5D; fig. S8B-8D).

We initially analyzed the Cnn1/Wip1-GFP complex in wild type and *cnn1-3A* lysates to ensure it assembles normally. The percentage of *CEN3* DNAs colocalizing with Wip1-GFP was indistinguishable between *dsn1-2D* versus *cnn1-3A dsn1-2D* lysates (54 ± 3% vs 58 ± 0.8 % mean ± S.D.) (Fig. 5C-5D, fig. S8B), consistent with our Cnn1 fluorescence microscopy results in cells (fig. S3B-3C). Next, we analyzed Ndc80c assembly. Nuf2-GFP colocalization with *CEN3* DNAs was also indistinguishable in *dsn1-2D* versus *cnn1-3A dsn1-2D* lysates (48 ± 1.1 % vs 48.8 ± 0.8 %) (Fig. 5C-5D, fig. S8C), indicating that the Cnn-3A mutant does not affect Ndc80c kinetochore recruitment. In contrast, Stu2-GFP colocalization showed a clear reduction in *cnn1-3A dsn1-2D* lysates compared to *dsn1-2D* (41 ± 1% vs 20 ± 2.3%) (Fig. 5C-5D, fig. S8D), consistent with immunoblots of Cnn1-3A purified material (Fig. 5A). These data suggest that the *cnn1-3A* mutation reduces Stu2 levels at the kinetochore without affecting Ndc80c levels.

To test whether Nuf2 and Stu2 levels are affected *in vivo* in the *cnn1-3A* mutant cells, we quantified Stu2 and Nuf2 localization to kinetochores (as previously described (*59, 60*)). WT and *cnn1-3A* cells containing Nuf2-GFP or Stu2-GFP were arrested in G1, released into the cell cycle, and harvested in metaphase. We used Ame1 tagged with the mKate fluorophore as an internal reference to identify kinetochore clusters. Consistent with the TIRF analysis, we observed that the relative Stu2 intensity at kinetochores in *cnn1-3A* mutant cells is reduced by ∼20% compared to wild type cells (Fig. 5E), whereas the Nuf2-GFP intensity is similar to wild type cells (fig. S8E). Taken together, our data suggests that mutating the C-region of Cnn1 reduces Stu2 levels at kinetochores both *in vivo* and *in vitro*.

### Cnn1 and Stu2 interact weakly *in vitro* independent of the C-region

Because the Cnn1 C-region promotes Stu2 association with kinetochores, we asked whether Cnn1 contacts Stu2 directly. We first used AlphaFold 3 multimer modeling to analyze the interaction between two full-length Stu2 molecules and a Cnn1 fragment (106-178 amino acids) that includes the C-region. The modeling did not predict an interaction with high confidence. However, it did suggest a lower confidence interaction between the dimerization interface of two Stu2 molecules with the Cnn1 C-region (iPTM =0.27 and pTM = 0.32) (fig. S9). To directly test this, we purified native Stu2 and Ndc80c from high-salt conditions to remove co-purifying proteins from budding yeast (*31*), and incubated each, alone or in combination, with immobilized recombinant wild type Cnn1 or Cnn1-3A (Fig. 6A, S10A, S10B). Ndc80c served as a positive control because it is the known Stu2 kinetochore receptor. As expected, we detected binding between Cnn1 and Ndc80c, and it was not affected by the presence or absence of Stu2. We also detected a weak interaction between full-length Cnn1 and Stu2, which was modestly enhanced when Ndc80c was present.

**Fig 6.**
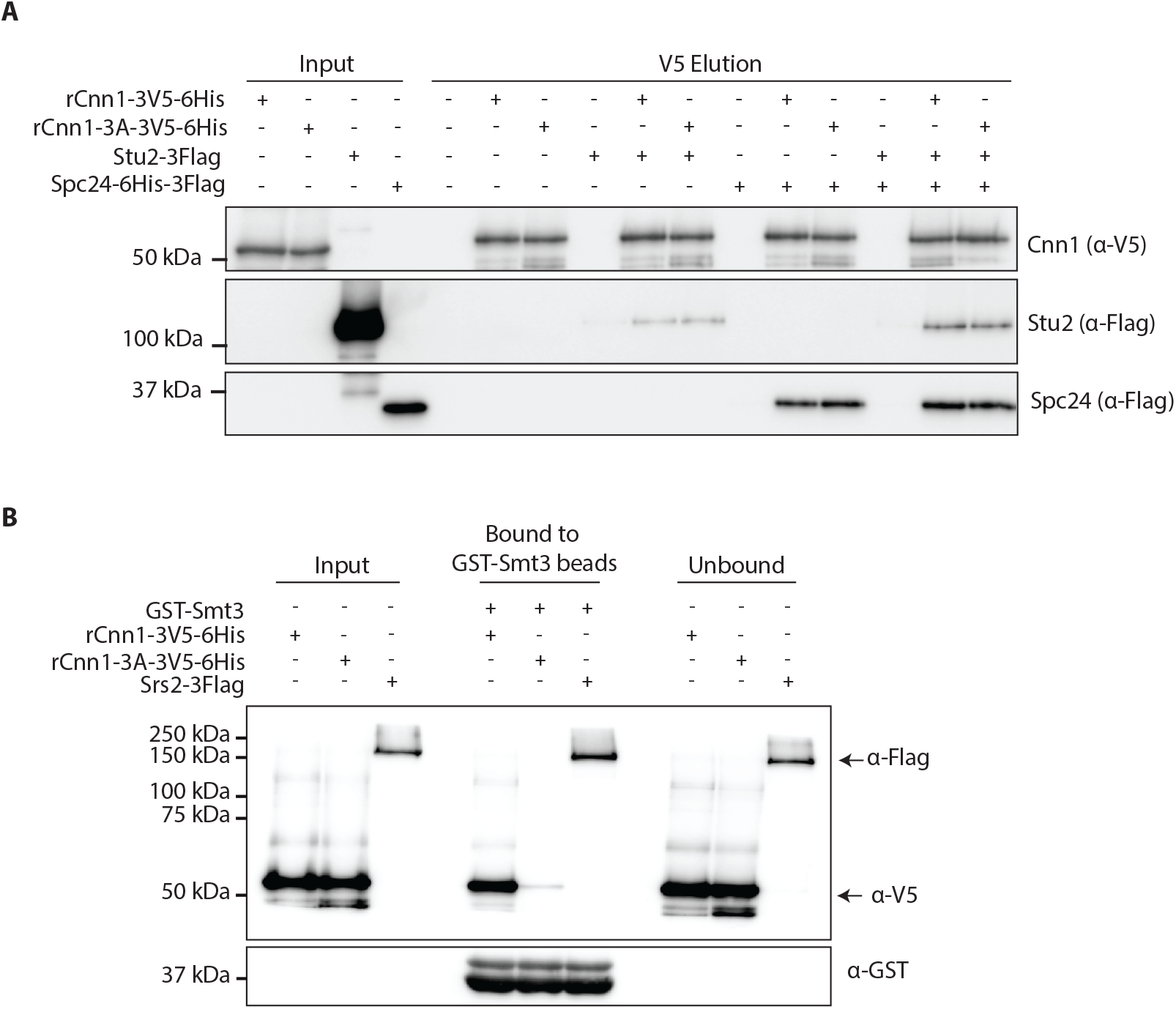
Cnn1 binds to the Stu2 and SUMO proteins. A. Binding assays between Cnn1 or Cnn1-3A and Stu2 and Ndc80c. Recombinant Cnn1 and Cnn1-3A with a 3V5 epitope tag at the C-terminus were immobilized on V5 beads that were subsequently incubated with an equal concentration of purified Stu2-Flag, Ndc80c (via Spc24-6His-3Flag) or both proteins for 30 mins at 4 °C before eluting with V5 peptide (1mg/ml). The proteins were analyzed via immunoblot. The input shows the protein added to the binding assay. B. Binding assay between Cnn1 or Cnn1-3A and Smt3. GST-Smt3 was immobilized on glutathione beads that were subsequently incubated with 2 μg of recombinant Cnn1-3V5-6His, recombinant Cnn1-3A-3V5-6His, or yeast purified Srs2-Flag for 4 hrs at 4 °C before eluting in sample buffer by boiling at 95 °C for 5 mins. Srs2-Flag served as a positive control for SUMO binding. The binding assays were analyzed via immunoblot. The input shows the purified proteins added to the binding assay.

However, the interaction between Stu2 and Cnn1 was not altered by the Cnn1-3A mutation, nor was the enhancement by Ndc80c affected (Fig. 6A). Thus, although Cnn1 and Stu2 can interact, this minimal interaction fails to recapitulate the C-region dependence observed in cells, in vitro kinetochore assembly, and laser trap assays. This discrepancy suggests additional features are required for the C-region to engage Stu2. Consistent with this possibility, a group-based prediction system (GPS) algorithm, SUMOsp, predicted that the C-region contains a SUMO-interacting motif (SIM), enabling it to bind sumoylated proteins (*61*). We therefore tested whether Cnn1 can bind to SUMO by immobilizing recombinant SUMO protein (GST-Smt3) (Fig 6B, S10C) on beads and incubating it with recombinant Cnn1 or Cnn1-3A. As a positive control, we also incubated the SUMO beads with the Srs2-3Flag protein purified from yeast (Fig. 6B, S10D) (*62*). As expected, we detected an interaction between Srs2-Flag and SUMO (Fig 6B). We also detected an interaction between Cnn1 and SUMO that was abolished by the Cnn1-3A mutations, suggesting that the Cnn1 C region is a bonafide SIM. It is therefore possible that sumoylation regulates the interaction between Cnn1 and Stu2 but was not present in our binding assays.

### Addition of Stu2 rescues the *in vitro* and *in vivo* defects associated with *cnn1-3A*

The observed reduction in Stu2 levels at Cnn1-3A kinetochores is consistent with the lower kinetochore-microtubule attachment strengths measured in Cnn1-3A purifications. Prior optical trapping assays established that kinetochore particles lacking Stu2 displayed significantly reduced rupture forces, and that adding Stu2 back rescued these weak attachments (*31*). We therefore tested whether the decreased strength of kinetochore microtubule attachments of Cnn1-3A *in vitro* can be rescued by adding Stu2. We purified Stu2-3Flag under high-salt conditions (Fig. S10A), incubated it with Cnn1 and Cnn1-3A particles for 30 mins at 4 °C, and then linked Cnn1 or Cnn1-3A to polystyrene beads for rupture force assays. We observed that Stu2 addition increases the attachment strength of the Cnn1-3A particles so that the survival curve nearly overlaps that of Cnn1 particles with *F*_1/2_ increasing from 4.8 pN (95% CI: 4.2 pN –5.2 pN) to 8.5 pN (95% CI: 7.6 pN –9.2 pN) (Fig. 7A). For the Cnn1 particles, the survival curves and *F*_1/2_ were not significantly changed by Stu2 addition (8.8 pN [95% CI: 8.1 pN – 9.7 pN]) vs 8.9 pN [95% CI:7.6 pN –10.9 pN]) (Fig. 7A). Mock treatment of Cnn1-3A particles with Stu2 storage buffer still resulted in weak kinetochore-microtubule interactions (4.3 pN [95% CI: 3.7 pN – 5.0 pN)). We also tested whether the addition of Stu2 that cannot interact with Ndc80 due to a deletion of its C-terminus (Stu2-(Δ855-888)) can restore the Cnn1-3A rupture forces (*30*). Cnn1-3A particles preincubated with Stu2-(Δ855-888) had an *F*_1/2_ of 5.1 pN (95% CI: 4.5 pN –5.9 pN). This was significantly lower than the *F*_1/2_ of 8.7 pN (95% CI: 7.3 pN –10.4 pN) for wild-type Stu2 and did not differ significantly from that of mock-treated Cnn1-3A particles (5.3 pN; 95% CI: 4.3 pN –6.1 pN) (Fig. 7B). Stu2 residues 855-888 are therefore required for Stu2 to strengthen attachments of Cnn1-3A particles, consistent with a role for Stu2 binding to the kinetochore.

**Fig 7.**
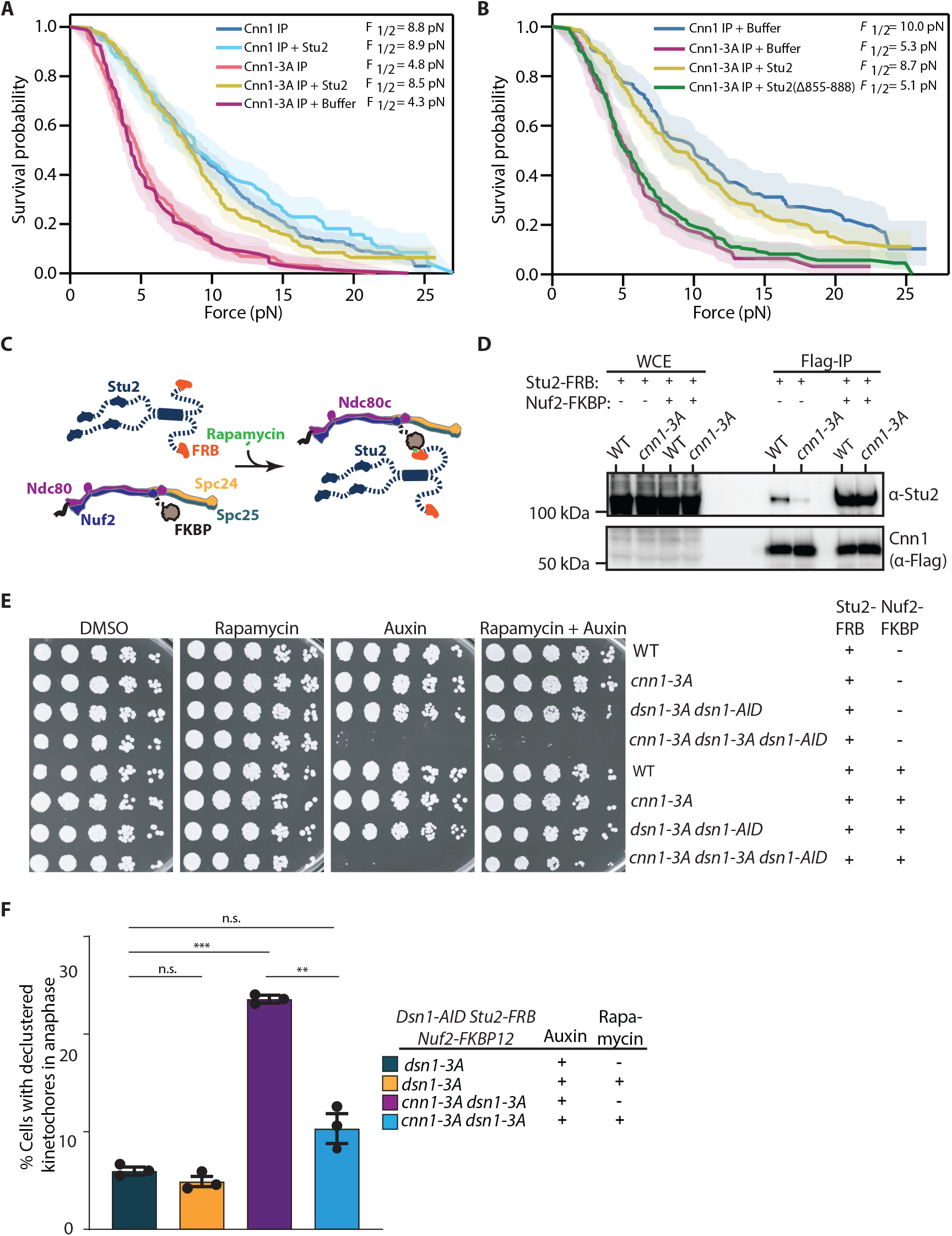
Addition of Stu2 rescues Cnn1-3A kinetochore-microtubule attachment defects. A. The Kaplan-Meier survival probability of bead attachment for rupture force experiments of wild type Cnn1 and Cnn1-3A particles (the previously mentioned rupture forces in Fig 4B were used for comparison) with and without purified Stu2 added back. Survival probability of Cnn1-3A particles with a buffer add back control were also performed. Shaded regions represent 95% confidence intervals (n = 378 for wild type Cnn1, n= 121 for wild type Cnn1 particles preincubated with Stu2, n= 175 for Cnn1-3A kinetochores, n = 210 for Cnn1-3A particles preincubated with Stu2, and n=91 for Cnn1-3A incubated with BH buffer) where n represents total events (rupture and censored). The p-value was calculated from the log-rank test (Cnn1 vs Cnn1+Stu2 p-value: 0.24; Cnn1 vs Cnn1-3A p-value: 2.0e-21; Cnn1-3A vs Cnn1-3A +Stu2 p-value: 1.2e-12; Cnn1 vs Cnn1-3A+Stu2 vs p-value 0.11: Cnn1-3A vs Cnn1-3A + Buffer p-value: 0.92; Cnn1 + Stu2 vs Cnn1-3A + Stu2: 0.03). B. The Kaplan-Meier survival probability of bead attachment for rupture force experiments of wild type Cnn1 and Cnn1-3A particles with addback of purified Stu2, Stu2(Δ855-888), or a mock treatment (Buffer H). Stu2 or Stu2(Δ855-888) were purified from cells depleted of Stu2-AID via an hour auxin treatment prior to harvesting. Shaded regions represent 95% confidence intervals (n = 94 for wild-type Cnn1 mock, n = 85 for Cnn1-3A kinetochores, n = 191 for Cnn1-3A particles preincubated with Stu2, and n = 210 for Cnn1-3A particles preincubated with Stu2(Δ855-888)) where n represents total events (rupture and censored). The p-value was calculated from the log-rank test: Cnn1 vs Cnn1-3A p-value: 3.7e-8; Cnn1 vs Cnn1-3A+Stu2 p-value: 0.21; Cnn1 vs Cnn1-3A+Stu2(Δ855-888) p-value: 4.6e-8; Cnn1-3A vs Cnn1-3A +Stu2 p-value 8.8e-8; Cnn1-3A vs Cnn1-3A +Stu2(Δ855-888) p-value 0.55; Cnn1-3A+Stu2 vs Cnn1-3A +Stu2(Δ855-888) p-value 1.7e-8. C. Schematic of the Stu2 tethering system to Ndc80c. Addition of rapamycin causes Stu2-FRB and Nuf2-FKBP to associate, restoring the interaction between Stu2 and kinetochores at the proper location (*30*). D. Cnn1-Flag and Cnn1-3A-Flag cells containing Stu2-FRB (SBY24829 and SBY24830) or Stu2-FRB Nuf2-FKBP (SBY24828 and SBY24831) were arrested in mitosis by benomyl treatment for 2.5 hours and 50 ng/ml rapamycin was added to induce the association between Stu2-FRB and Nuf2-FKBP one hour prior to harvesting. Cnn1-Flag was immunoprecipitated and Stu2 recruitment was analyzed by immunoblot (n=2). E. Five-fold serial dilutions of wild type (SBY24829, SBY24828), *cnn1-3A* (SBY24831, SBY24830), *dsn1-AID dsn1-3A* (SBY24900, SBY24833), *cnn1-3A dsn1-AID dsn1-3A* (SBY24902, SBY24835) cells expressing Stu2-FRB with or without Nuf2-FKBP were assessed by spotting cells on YPD plates containing DMSO, rapamycin, auxin, or rapamycin and auxin together. Cells were incubated at 23 °C and images were taken 72 hours after plating. F. *Dsn1-3A STU2-FRB NUF2-FKBP* (SBY25566) and *cnn1-3A dsn1-3A STU2-FRB NUF2-FKBP* (SBY25569) cells were arrested in G1 with alpha factor for three hours and auxin was added for the last 30 minutes. The cells were then released into YPD containing auxin with or without rapamycin. Cells were harvested 75 and 80 minutes after release during metaphase to analyze kinetochores. The percentage of cells with declustered kinetochore-microtubule attachments in each strain during metaphase was quantified across 3 experiments (n > 200 total cells per strain). The error bar represents standard error. The p-value was determined using Welch’s t-test. The p-value between *dsn1-3A STU2-FRB NUF2-FKBP* with or without rapamycin is 0.2, *dsn1-3A STU2-FRB NUF2-FKBP* and *cnn1-3A dsn1-3A STU2-FRB NUF2-FKBP* in auxin media is 0.009, *cnn1-3A dsn1-3A STU2-FRB NUF2-FKBP* with or without rapamycin is 0.001 and *dsn1-3A STU2-FRB NUF2-FKBP* and *cnn1-3A dsn1-3A STU2-FRB NUF2-FKBP* in auxin media with or without rapamycin is 0.09. ** represents <0.01. n.s. represents non-significant.

We next asked whether restoring Stu2 to kinetochores in *cnn1-3A* cells is sufficient to rescue kinetochore function. Because Stu2 overexpression alters spindle dynamics, it was not a good way to test for rescue of the growth defects of the *cnn1-3A dsn1-3A* cells (*63*). However, Stu2 tethered to Nuf2 via an FRB-FKBP rapamycin-induced dimerization system is functional at kinetochores because it restores Stu2 to its proper position in the kinetochore (Fig. 7C) (*30*). We therefore asked whether introducing Stu2-FRB and Nuf2-FKBP into *cnn1-3A dsn1-3A* cells rescued their growth defect. The endogenous Stu2 and Nuf2 proteins were tagged, and the strains also carried a mutant *TOR1* (*TOR1-1*) and *fpr1Δ*, rendering them rapamycin-resistant. We first tested whether tethering Stu2 to Nuf2 increases its association with Cnn1 purifications. We added rapamycin to wild type and *cnn1-3A* cells containing *STU2-FRB* with or without *NUF2-FKBP* and arrested them in mitosis by benomyl. We purified Cnn1 via the Flag tag at its C-terminus and detected significant recruitment of Stu2 to the Cnn1 and Cnn1-3A purifications in the presence of Nuf2-FKBP, indicating the tethering system worked well (Fig. 7D).

To test whether Stu2 tethering rescues the growth defect of the *cnn1-3A dsn1-3A* cells, we introduced Stu2-FRB in the presence or absence of Nuf2-FKBP into wild type, *cnn1-3A, dsn1-3A* and *cnn1-3A dsn1-3A* strains and plated them on DMSO, auxin, rapamycin, or both drugs together (Fig. 7E). We observed lethality in *cnn1-3A dsn1-3A* cells on plates containing auxin with or without rapamycin. The phenotype of the *cnn1-3A dsn1-3A* double mutants was more severe in combination with Stu2-FRB and Nuf2-FKBP than what was observed without these tags in figure 2B. Strikingly, the viability of these *cnn1-3A dsn1-3A* cells was completely rescued when Stu2 was artificially tethered to the kinetochore. Consistent with this, we analyzed kinetochore clustering and found that tethering Stu2 also suppressed the declustered kinetochores we observed in *cnn1-3A dsn1-3A* cells (Fig 7F). We note that the tethering configuration brings more Stu2 to kinetochores than is normally present, so it is possible that this high level overcomes another function for the C-region. However, these data are most consistent with the possibility that the Cnn1 C-region contributes to the association and function of Stu2 at kinetochores.

## DISCUSSION

Here we show that Cnn1-based kinetochore particles can maintain load-bearing attachments to dynamic microtubule tips in vitro and track microtubule tips under force, similar to Dsn1-based particles (*42*). We further demonstrate that a conserved region in the N-terminal tail of Cnn1, the C-region, is critical for maintaining kinetochore-microtubule coupling. Mutations of the C-region lead to kinetochore de-clustering and activation of the spindle assembly checkpoint in *dsn1-3A* cells. As discussed below, the underlying defect in the Cnn1 mutant appears to be a perturbed Stu2 interaction with the kinetochore that leads to weakened kinetochore-microtubule attachments (*31*).

We assayed the biophysical properties of the Cnn1 particles using an optical trapping assay in which the Cnn1 or Dsn1 proteins were directly tethered to the trapped beads, so force was transmitted through each receptor. The comparable rupture forces we measured for Cnn1-versus Dsn1-purified particles suggests similar microtubule-attachment architectures, although the interdependent recruitment of the two receptors limits the conclusion (*58*). As previously demonstrated for Dsn1-purified kinetochore particles, Cnn1-purified particles support significantly higher forces than individual Ndc80 complexes, suggesting the Cnn1 particles carry multiple Ndc80 complexes and additional outer kinetochore proteins that contribute to their coupling strength(*64*). Intriguingly, although Cnn1 binds Ndc80c in a 1:1 stoichiometry via its conserved N-terminal helical domain, prior work suggests that the human CENP-T protein might help to cluster multiple Ndc80 complexes, which could contribute to its ability to support strong microtubule attachments (*19, 65, 66*). In the future, it will be important to compare additional properties of the receptors, such as the length of time they can maintain attachments to dynamic microtubule tips under various forces.

Our work also found that the C-region of Cnn1 is critical for sustaining kinetochore-microtubule attachments. Mutations in this region significantly weaken the attachment between Cnn1-based particles and dynamic microtubules, with rupture forces similar to Dsn1-based particles lacking Stu2 (*31*). Although recent cross-linking mass spectrometry suggests the C-region might be a contact point for Ndc80c, we did not detect reduced Ndc80c levels in immunoblots of mutant C-region purifications (*46*). Likewise, our single-molecule assembly assay did not detect reduced Ndc80c levels, suggesting the functional defect in the C-region mutant is not caused by decreased Ndc80c.

We found that the Cnn1 C-region alters Stu2 levels even though Ndc80c is the only known kinetochore receptor for Stu2. Consistent with this, we found that the Stu2 co-purification with Cnn1 required Cnn1’s interaction with Ndc80. However, the Stu2-Ndc80 interaction is weak (kD=48 micromolar) (*30*), so Stu2 might require additional factors to stably localize to kinetochores in cells. Stu2 is sub-stoichiometric to Ndc80c at kinetochores and the Stu2-Ndc80 contact alone is not sufficient for kinetochore localization *in vivo*, which also requires Stu2 dimerization and the TOG1 domain (*31*). We therefore considered the possibility that a second, transient contact with Cnn1 stabilizes Stu2 on Ndc80c. However, although we detected a weak interaction between Cnn1 and Stu2 *in vitro*, it was not affected by the Cnn1 C-region mutation.

One possible explanation is that our in vitro interaction assay lacked post-translational modifications needed for the interaction. Consistent with this, we found that the C-region binds to SUMO and Stu2 is known to be sumoylated, so it is conceivable that sumoylation regulates the interaction but was missing in the Stu2 protein purification we used (*61, 67*). Another possibility is that the Cnn1 kinetochore receptor (Ctf3 complex) needs to be present to position Cnn1 in a manner that could affect Stu2 association. In the future, additional protein interaction experiments will be needed to determine whether Cnn1 and Stu2 directly interact and how this potential interaction is regulated by the C-region and sumoylation. It will also be interesting to determine whether CENP-T affects chTOG kinetochore association in other species since CENP-T contains SUMO-interacting motifs similar to the Cnn1 C-region and the Cnn1 and CENP-T complexes share a conserved architecture (*18*).

In summary, our work provides insight into the role of Cnn1 at the kinetochore. Cnn1 purified particles support much higher forces than Ndc80 complexes, similar to Dsn1 purifications ((*64*)). Our data suggests that the Cnn1 C-region contributes to this coupling strength by promoting Stu2 localization rather than through recruiting Ndc80. The dramatic effect of partial loss of kinetochore Stu2 on microtubule coupling strength is consistent with Stu2 acting to stabilize multiple Ndc80 complexes. One possibility is that Stu2 dimers bridge pairs of Ndc80 complexes, which facilitate Ndc80c clustering and enhances the strength of kinetochore-microtubule attachments (*66*). Dsn1 also recruits Ndc80c, so it is important to determine whether Stu2 interacts with Dsn1 to strengthen kinetochore-microtubule attachments. More broadly, future studies will continue to elucidate the underlying mechanisms that ensure kinetochores make accurate load-bearing attachments to microtubules.

## MATERIALS AND METHODS

### Strain construction and microbial techniques

Standard yeast genetic crosses, media preparation, and microbial techniques were used (*68*). *Saccharomyces cerevisiae* strains used in this study are derivatives of the W303 genetic background (SBY3) and are listed in Table S1. Epitope-tagged versions of Cnn1 (6His-3Flag, 3Flag or 3mNeonGreen) were constructed by standard PCR-based integration as described (*69*). The *cnn1-3A* and *cnn1(Δ157-163)* mutations were made using a single-step CRISPR/Cas9 edit as previously described (*70*). The plasmid targeting the CRISPR edit was co-transformed with the repair templates SB7634 (*cnn1-3A*) and SB8713 (*cnn1(Δ157-163)*). The mutations were confirmed by PCR followed by sequencing. To generate strains expressing ectopic *dsn1-3A* at the *HIS3* locus, PCR was performed on the plasmid pSB1108 (described in (*71*)). The PCR product was digested with NheI, followed by transformation into cells expressing endogenous *DSN1-3HA-IAA7* (construction of *3HA-IAA7* tagging plasmid was described previously in (*31*)) and the *pGPD1-TIR1* allele integrated at the *LEU2* locus (described in (*41*)). Construction of yeast strain containing *DSN1-6His-3Flag* is described in (*42*), *SPC24-6His-3Flag* in (*31*), *GFP-TUB1* in (*72*), *PDS1-13Myc* in (*73*), *STU2-3Flag* (*31*), *STU2-GFP* in (*58*), *AME1-mKATE* in (*74*), *mad3Δ* in (*75*)*, NDC10-mCherry* (*43*). Yeast expressing *MTW1-mCherry* were a kind gift from Trisha Davis’ laboratory (University of Washington) and yeast expressing *STU2-FRB* and *NUF2-FKBP* were a kind gift from Matthew Miller’s laboratory (University of Utah) and were described in (*30*).

Yeast strains for ectopic expression of Stu2(Δ855-888)-3Flag (SBY25588) and Stu2-3Flag (SBY25592) were constructed by digesting pSB3779 and pSB3780 with SwaI and transforming into yeast. The integrated sequence was confirmed by sequencing PCR of genomic DNA with primers SB4819/ SB4820 and protein expression and epitope tagging were confirmed by immunoblot. Final strains were generated by crossing with cells expressing endogenous *STU2-3HA-IAA7* and *pGPD1-OsTIR1* allele integrated at the *HIS3* locus. Construction of *3HA-IAA7* tagging plasmid was described previously in (31)

### Plasmid construction

To generate plasmids to make CRISPR edits on *CNN1* to create *cnn1-3A* and *cnn1(Δ157-163)*, the parental plasmid, pSB3218 (a gift from Elçin Ünal’s laboratory; University of Berkeley, CA), was linearized by digesting with BsmBI and assembled with guide RNAs (*cnn1-3A* (SB7632 and SB7633) and *cnn1(Δ157-163)* (SB8711 and SB8712)) by the Gibson method (*76*) resulting in *cnn1-3A* (pSB3405) and *cnn1(Δ157-163)* (pSB3740) plasmids.

To generate a recombinant construct of *CNN1-3V5-6His* or *cnn1-3A-3V5-6His*, plasmid pSB3524 (described in (*77*)) was amplified using primers SB8883 and SB8884, and *CNN1* or *cnn1-3A* was amplified from gDNA of strains SBY24005 and SBY24006, respectively, using primers SB8885 and SB8886. The PCR products were assembled by the Gibson method, generating plasmids pSB3667 and pSB3741. The plasmids have pET-21 b (+) backbone (Novagen). All the plasmids used are listed in Table S2. Primers used to construct the strains and plasmids are listed in Table S3. Strains and plasmids used in this study are available upon request.

Single integration *LEU2* vectors for *STU2-3Flag* (pSB3780) and *stu2(Δ855-888)-3Flag* (pSB3779) were prepared through common cloning techniques from single-copy *LEU2* integration plasmids pSB2232 (*STU2(WT)-3V5*) and pSB2358 (*stu2(Δ855-888-3V5*)(*30*). Briefly, pSB3780 was made by replacement of the 3V5 epitope tags with 3Flag by amplifying pSB2232 with oligos SB9567/ SB9568 and pSB812 with oligos SB9565/ SB9566 and performing Gibson assembly. pSB3779 was prepared the same way by amplifying pSB2358 with oligos SB9563/ SB9564 and pSB812 with oligos SB9561/ SB9562. The resulting plasmids were verified with whole plasmid sequencing.

### Serial dilution cell viability assay

*S. cerevisiae* strains were grown overnight in YPD medium. Cell densities were measured using a spectrophotometer, and cultures were diluted with sterile water in a 96-well plate to normalize densities across strains for the initial well. The cultures were then serially diluted 5-fold using sterile water. Cells from each well were then spotted onto YPD plates with DMSO, 500 μM Auxin, 50 ng/ml Rapamycin, or both 500 μM Auxin and 50 ng/ml Rapamycin. The plates were incubated for 1-3 days at 23 °C and imaged after sufficient growth.

### Auxin-inducible degradation

The auxin inducible system was previously described (*78*). Briefly, auxin-induced degradation of *dsn1-AID* cells expressing *DSN1-3HA-IAA7*, in the presence of *pGPD1-TIR1,* was performed by treating cells with 500 μΜ IAA (indole-3-acetic acid dissolved in DMSO) for 1 hour before harvesting or imaging. Similarly, for purification of ectopic Stu2 or Stu2(Δ855-888), auxin-induced degradation of endogenous Stu2 in cells expressing *STU2-3HA-IAA7*, in the presence of *pGPD1-OsTIR1,* was performed by treating cells with 500 μΜ IAA for 1 hour before harvesting.

### FRB/FKBP tethering

For tethering Stu2-FRB with Nuf2-FKBP in culture, 50 ng/mL (55 nM) rapamycin was added 1 hour prior to harvesting (*30*). The cell viability assays were carried out as indicated above, except that overnight cultures were diluted and grown for several hours to OD∼1.0-2.0 before the dilution with water to normalize cell numbers. Rapamycin was added to YPD agar plates at a final concentration of 50 ng/ml, with or without 500 μM auxin.

### Fluorescent Microscopy and Quantification

Cells expressing fluorescently tagged kinetochore proteins were arrested in G1 using 1 μg/ml alpha factor. After washing three times with YPD, cells were resuspended in YPD media containing either auxin (to degrade the endogenous Dsn1-AID when indicated) or DMSO (control). 1 ml cells were collected at 75 minutes after G1 release to visualize the metaphase stage (for Cnn1 localization, cells were also collected at 100 minutes for the anaphase stage) and washed once with sterile water. The cell cycle stage was determined based on spindle length.

Cells with short spindles were classified as metaphase and long spindles as anaphase. Next, the cells were fixed with 3.7% formaldehyde in 0.1 M phosphate buffer (pH 7.5) at room temperature for 5 minutes. Fixed cells were washed with water once and resuspended in the same buffer containing 1.2 M Sorbitol. Cells were then mounted on 1.5% agar pads (made in 0.1 M phosphate buffer, pH 7.5) and sealed with liquid VALAP (1:1:1 petroleum jelly:lanolin:paraffin wax).

Cells were imaged on a Deltavision Ultra deconvolution high-resolution microscope equipped with a 100×/1.4 PlanApo N oil-immersion objective (Olympus). Z-stacks were acquired across the entire cell at 0.5 µM increments. Images were deconvolved in softWorX v7.2.1 (GE) using standard settings. Projections were made using a maximum-intensity algorithm. Composite images were assembled, and false coloring was applied with Fiji.

To quantify kinetochore protein fluorescence intensity, maximum intensity projections of the sister kinetochores were circled in Fiji using the freehand tool, and the total signal intensity of each kinetochore was obtained. Background intensity was subtracted from the total intensity to obtain the corrected signal intensity of the kinetochore pair, which was then summed up for each cell. Next, the corrected fluorescent intensity of the kinetochore protein was divided by the corrected signal of the fiducial kinetochore marker to get the normalized fluorescence intensity.

### Protein Biochemistry

#### Native Dsn1 and Cnn1 particle purifications

Native Dsn1 and Cnn1 particles were purified from cells arrested in mitosis by benomyl addition (30 μg/ml for 2.5 hours). Purifications were performed as described previously (*42*). Briefly, cells were grown to late log phase OD_600_ = 2-3 and harvested by centrifugation at 5,000 g for 5 minutes. For experiments testing Cnn1 particle binding to microtubules in the presence or absence of the *ndc80-1* allele, benomyl was added to cultures of SBY24005 and SBY24285 and shifted to 37C for 3 hrs prior to harvesting at OD600 = 2-3. Next, the cell pellets were washed once in cold water containing 0.2 mM PMSF (10-17.5 ml of water + PMSF for every 500 ml culture harvested) and then in lysis buffer: Buffer H (BH) 0.15 (25 mM Hepes pH 8, 2 mM MgCl2, 0.1 M EDTA, 0.5 M EGTA, 15% glycerol, 0.1% NP-40, 150 mM KCl) supplemented with phosphatase inhibitors (1 mM sodium pyrophosphate, 2 mM sodium-β-glycerophosphate, 0.1 mM sodium orthovanadate, 5 mM sodium fluoride, and 0.2 μM microcystin-LR) and protease inhibitors (0.2 mM PMSF, 10-20 μg/ml leupeptin, 10-20 μg/ml pepstatin, 10-20 μg/ml chymostatin). Cells were pelleted at 3,000 g for 5 mins, resuspended in lysis buffer in a volume determined using the following calculation: OD x ml culture x 1.25. Resuspended cells were flash frozen in liquid nitrogen and lysed in a freezer mill with 10 cycles of 2 minutes on, 2 minutes off (SPEX SamplePrep). The lysate powder was thawed and incubated with benzonase (Millipore Sigma 71205-3, 50 units/ml lysate) for 30 minutes at 4 °C with constant rotation. Next, the lysate was ultracentrifuged at 98,500 g for 90 mins at 4 °C. Protein concentration was assessed using a Pierce BCA Protein Assay Kit (Thermo Scientific) and normalized with lysis buffer prior to IP. Protein G Dynabeads (ThermoFisher Scientific) were crosslinked with an α-M3DK antibody (using dimethyl pimelimidate) that recognizes the 3-Flag epitope tag (Genscript)(*79*). Beads were added to the lysate (15 μl beads per 12.6 mg protein) and incubated at 4 °C for 3 hours with constant rotation. Next, the beads were washed 3 times with lysis buffer containing 2 mM DTT, protease and phosphatase inhibitors, and twice with the same buffer without DTT. The protein was eluted in BH0.15 buffer containing 1 mg/ml Flag peptide (Sigma-Aldrich) in 1/3 bead volume by gentle agitation for 30 minutes at 23 °C.

#### High salt purification of Stu2 and Ndc80

Cells expressing Stu2-3Flag (SBY12275) or Stu2-3V5 (SBY11709) were used to purify Stu2. The cells were lysed and resuspended in lysis buffer: Buffer H (BH) 1.0 (25 mM Hepes pH 8, 2 mM MgCl2, 0.1 M EDTA, 0.5 M EGTA, 15% glycerol, 0.1% NP-40, 1M KCl). To purify Stu2-3Flag (SBY25592) and Stu2(Δ855-888)-3Flag (SBY25588), cells were treated with benomyl for 90 minutes before 0.5 mM IAA was added for 1 hour prior to harvest (2.5 hour total benomyl treatment). Like other Stu2 purifications, cells were washed and lysed in high salt lysis buffer: Buffer H (BH) 1.0 (25 mM Hepes pH 8, 2 mM MgCl2, 0.1 M EDTA, 0.5 M EGTA, 15% glycerol, 0.1% NP-40, 1M KCl). For Ndc80c purification, cells expressing Spc24-6His-3Flag and Spc105-AID (SBY14022) were used. The cells were treated with 500 μM auxin 45 minutes prior to harvesting to prevent co-purification of Spc105-Kre28. Lysed cells were resuspended in BH, as described above but with 750 mM KCl instead of 1 M. For Srs2, cells expressing Srs2-3Flag (SBY25556) were grown and BH/0.40 (400 mM KCl) was used as lysis buffer. Dynabeads conjugated to α-Flag were incubated with lysate for 3 hr with constant rotation, followed by three washes with BH containing protease and phosphatase inhibitors, 2 mM dithiothreitol (DTT), and 1 M KCl. Beads were then further washed twice with BH containing 150 mM KCl with protease and phosphatase inhibitors. For purification of Stu2 and Stu2(Δ855-888), phosphatase inhibitors were omitted from the wash steps following the immunoprecipitation. The proteins were then eluted from the beads by gentle agitation in elution buffer (1 mg/ml 3Flag peptide in BH with 150 mM KCl) for 30 mins at room temperature, using half the bead volume. The concentrations of Stu2, Ndc80c, and Srs2 were assessed by SDS-PAGE relative to a BSA standard curve.

#### Recombinant protein purification

To purify recombinant Cnn1, pSB3667 (*CNN1-3V5-6His*) and pSB3741 (*cnn1-3A-3V5-6His*) were transformed into Rosetta 2 (DE3) pLysS cells (Novagen). 1 L cultures were grown in Terrific Broth (12 g/L tryptone, 24 g/l yeast extract, 0.4% v/v glycerol, 0.017 M KH_2_PO_4_, 0.072 M K_2_HPO_4_) supplemented with 25 μg/ml ampicillin. Once the OD_600_ reached 0.4, 0.5 mM IPTG was added to induce protein production for 3 hours at 37 °C. Cells were harvested by centrifugation at 5000 g for 20 minutes, and the cell pellets were washed with water containing 0.2 mM PMSF. Next, the cells were resuspended in sonication buffer containing B/ 0.3: 25 mM Hepes pH 8, 2 mM MgCl2, 15% glycerol, 0.1% NP-40, 300 mM KCl, along with 0.2 mM PMSF, protease inhibitor (1 tablet of cOmplete Protease inhibitor cocktail, EDTA-free; Sigma-Aldrich; for 30ml of buffer) and 10 mM imidazole. The resuspended cells were sonicated for 5 minutes at 50% amplitude with 1 min on and 1 min off. The extract was spun at 11,000 x g for 30 min to pellet debris, and the clarified lysate was incubated with equilibrated HisPur Cobalt resin (Thermo Fisher Scientific) on a rotary mixer for 30 min at 4 °C. Resin was collected in a gravity column (column used had 2 ml column volume). The resin was washed 6 times with 2 column volumes with the wash buffer (BH/0.3 + 0.2 mM PMSF + 10 mM Imidazole). Protein was eluted with BH/0.3 + PMSF + 150 mM imidazole. 6 eluates were collected, each of 1 column volume. The purified eluates were run on SDS-PAGE for verification. The second eluate was used for the *in vitro* binding assay (fig. S10B). Protein concentration was assessed by SDS-PAGE using a BSA standard curve.

To purify recombinant Smt3, pSB3776 (*GST-SMT3*) was transformed into NiCo21 (DE3) competent cells (NEB). A 1L culture was grown in 2XTY (16 g/L tryptone, 10 g/L yeast extract, 5 g/L NaCl) supplemented with 100 μg/ml ampicillin. Protein expression was induced with 0.2 mM IPTG overnight at 16 °C. Cells were lysed via sonication in 1X PBS, 1M NaCl, 0.05% Tween-20, 1 mM PMSF, 10 mM DTT and the lysate was spun at 15,000 x g for 1 hour at 4C. Lysate was loaded onto a column of Glutathione Sepharose 4B resin (Cytiva). The column was washed with 100 mL wash buffer (1X PBS, 0.05% Tween-20, 0.5 mM DTT, 0.25M KCl), followed by 25 mL wash buffer lacking Tween-20. Protein was eluted with 5 mM glutathione, 50 mM HEPES pH 8, 100 mM KCl and dialyzed overnight into 50 mM HEPES pH 8, 150 mM KCl, 30% glycerol. Protein purity and concentration was assessed by Coomassie staining of a SDS-PAGE gel using a BSA standard curve.

#### In vitro binding assay

Purified Cnn1-3V5-6His (∼700 ng) or Cnn1-3A-3V5-6His (∼600 ng) were immobilized on α-V5 conjugated Protein G dynabeads (ThermoFisher Scientific), by incubating the recombinant protein for 2 hours with constant rotation at 4 °C. Beads were washed 4 times with Buffer H 0.15 supplemented with protease inhibitors (0.2 mM PMSF, 20 μg/ml leupeptin, 20 μg/ml pepstatin, 20 μg/ml chymostatin) and were split into four aliquots to perform binding reactions between Cnn1 or Cnn1-3A with: 1) Buffer H/0.15, 2) with purified Stu2 (∼550 ng), 3) with purified Ndc80c (∼300 ng), 4) with both Stu2 and Ndc80c together. Beads with the respective proteins for the binding assays were rotated for 30 minutes at 4 °C. After the incubation, beads were washed twice with BH/0.15 supplemented with protease and phosphatase inhibitors. Bound proteins were eluted from the beads by gentle agitation with elution buffer (1 mg/ml V5 peptide in BH 0.15). V5-elution was loaded onto a 10% SDS-PAGE gel, and binding was analyzed by immunoblotting (Fig. S7C).

For the Smt3 binding assay, purified recombinant GST-Smt3 (SUMO mature form SMTgg) was bound to PureCube Glutathione MagBeads (Cube Biotech) in BH/0.05 (50 mM KCl) for 1 hour with constant rotation at 4 °C (*80*). The beads were washed 3 times in BH/0.05 and then incubated with ∼2 g purified recombinant Cnn1-3V5-6His, Cnn1-3A-3V5-6His, or yeast natively purified Srs2-3Flag in BH/0.05 for 4 hours at 4 °C. Supernatant (unbound) was taken, the beads were washed 3 times in BH/0.05, and bound protein was eluted by boiling beads in sample buffer. Input, bound, and supernatant samples were analyzed via immunoblot.

#### Immunoblot and silver stain gel analysis

For immunoblot analysis, cell lysates were prepared either by bead-beating with acid-washed glass beads in SDS (sodium dodecyl sulfate) buffer or as described above for Dsn1 and Cnn1 protein purifications. Protein samples were separated by 7.5-10% SDS-PAGE and transferred to a 0.45 μm nitrocellulose membrane using a wet transfer apparatus (Hoefer). Commercial antibodies used for immunoblotting were as follows: α-Pgk1 (Invitrogen; 4592560; 1:10,000), α-mNeonGreen (Cell signaling technology; E8E3V; 1:1,000), α-Μyc (Cell signaling technology; 71D10; 1:1,000), α-M3DK (to recognize Flag epitopes; GenScript; 1:10,000), α-V5 (Invitrogen; R96025; 1:5,000), α-Cse4 (GenScript; 9536; 1:500), α-Ctf19 (GenScript; 1:5,000), α-Spc105 (GenScript; 1:10,000), α-Ask1 (GenScript; 1:3,000, previously described in (*45*)). α-Ndc10 (OD1) used at 1:5,000 dilution and α-Ndc80 (OD4) used at a 1:10,000 dilution were kind gifts from Arshad Desai. The α-Stu2 (1:1,000) and α-Spc34 (1:5,000) antibodies were affinity-purified from polyclonal Stu2 or Spc34 sera (raised in rabbit using recombinant full-length proteins) custom generated by GenScript. The α-Spc34 antibody was verified by immunoblot of whole cell lysates comparing the band shift between Spc34 and Spc34-YFP. The α-Stu2 antibody was similarly verified by comparing Stu2 to Stu2-13Myc on an immunoblot. Secondary antibodies used were sheep anti-mouse antibody conjugated to HRP (GE Biosciences) at 1:10,000 or donkey anti-rabbit antibody conjugated to HRP (GE Biosciences) at 1:10,000. Antibodies were detected using SuperSignal West Dura Chemiluminescent Substrate or SuperSignal West Femto Substrate (Thermo Fisher Scientific). Immunoblots were imaged with a ChemiDoc MP system (Bio-Rad). For silver staining analysis, protein samples were separated using a precast 4-12% Bis-tris gel (Thermo Fischer Scientific) using MES buffer and stained using the Silver Quest Staining Kit (Invitrogen).

#### Mass spectrometry

The Flag tag on Dsn1-6His-3Flag or Cnn1-6His-3Flag was used to purify particles, as described previously (in the “native kinetochore particle purification” section). 10 μl of the eluate was run on a 4-12% Bis-Tris gel and analyzed by silver staining. The remaining eluate was precipitated overnight at 4 °C with 20% TCA. The next day, the precipitated particles were centrifuged at 16,000 g for 30 minutes at 4 °C. The supernatant was discarded. The pellet was washed three times with 1M HCl-acetone, air-dried at room temperature, resuspended in 10 μl sample buffer, and boiled at 95 °C for 5 minutes. Each sample was run on a 4-12% Bis-Tris gel (Thermo Fisher Scientific) for 5 minutes at 100V. The gel was stained overnight with 0.1% Coomassie G-250 and destained the following day. The gel section above the 3Flag peptide band was excised. The excised gel was subjected to in-gel digestion with Trypsin for LC-MS/MS analysis (*81*). The extracted peptide sample was acidified using 30% formic acid at room temperature for 1 hour.

Samples were then spun down, and the supernatant was collected. Samples were desalted using ZipTipC18 (Millipore) and eluted with 70% acetonitrile/0.1% trifluoroacetic acid. The desalted samples were concentrated using a speed vacuum. The resulting peptide samples were analyzed by LC-MS/MS using a Thermo Fisher Scientific Easy-nLC 1000 coupled to a Thermo Fisher Scientific Orbitrap Fusion mass spectrometer. Thermo Fisher Scientific Proteome Discoverer v3.1, with Sequest HT as the protein database search algorithm, was used to analyze the mass spectrometry data. The data were searched against the SGD yeast (W303_JRIU00000000_SGD_060922) database, which included common contaminants. A tryptic enzyme constraint and allowance for up to 2 missed cleavages was used. The precursor ion tolerance was set to 10 ppm, and the fragment ion tolerance was set to 0.6 Da. Variable modifications included oxidation on methionine (+15.995 Da) and phosphorylation on serine, threonine, and tyrosine (+79.966 Da). Static modifications included carbamidomethyl on cysteine (+57.021 Da). Peptide validation was performed using Percolator. Peptide identifications were filtered to a 1% false discovery rate.

For Supplemental Fig. 1A, the raw PSM counts for each kinetochore protein purified from IP of Dsn1-6His-3Flag or Cnn1-6His-3Flag are presented. The raw data is deposited at: https://massive.ucsd.edu/ProteoSAFe/private-dataset.jsp?task=8ce17ed601e343ae9edfcfbfa7f2fe92.

### Cell cycle progression and chromosome segregation assay

Exponentially growing *S*. *cerevisiae* cells were arrested in G1 using 1 μg/ml α-factor for 3 hours at 23 °C. 30 minutes prior to washing away the pheromone, 500 μM Auxin was added to degrade endogenous Dsn1-AID. The cells were then washed three times and resuspended in the YPD media containing 500 μM Auxin. 1 ml of cells was collected every 20 minutes for 180 minutes, and α-factor was re-added after 70 minutes to prevent entry into the subsequent cell cycle.

For budding index determination, cells at every time point were fixed using 3.7% formaldehyde. The cells were visualized under a Nikon Eclipse E200 microscope equipped with a 40X objective (Nikon) to calculate the budding index.

For Pds1 time courses, OD_600_ of the cultures was measured at every time point, and 1 ml of cells was harvested. Laemmli buffer was added to the pellets (volume calculated using culture OD_600_ as a reference to normalize protein concentrations), and lysate was prepared by bead-beating (Biospec Products) using 0.5 mm glass beads. Samples were boiled for 5 min at 95 °C, and 10 μL was loaded onto a 10% SDS-PAGE gel, followed by immunoblotting. Pgk1 was used as the loading control.

For the chromosome segregation assay (*55*), Chromosome 8 was fluorescently tagged by integrating a tandem array of LacO sequence, 2.1 kb from *CEN8* (a kind gift from Marion A. Shonn described in (*82*)) in cells expressing a LacI-GFP fusion protein. The cells were fixed with 3.7% formaldehyde at 100 minutes after G1 release, and the segregated chromosomes were visualized by the Deltavision Ultra deconvolution high-resolution microscope equipped with a 100×/1.4 PlanApo N oil-immersion objective (Olympus). DAPI staining was used to determine the cell cycle stages.

### TIRFM imaging and analysis

TIRFM was performed as previously described (*57, 58*). Briefly, all images were acquired with a Nikon TE2000 inverted RING-TIRF microscope at a resolution of 512 × 512 pixels, a pixel size of 0.11 μm/pixel, and a readout speed of 10 MHz. Atto-647-labeled *CEN3* DNAs were excited at 640 nm for 300 ms, GFP-tagged proteins at 488 nm for 200 ms. Images were analyzed with CellProfiler (4.2.6) to determine colocalization and quantify signal overlap between the DNA channel (647 nm) and the GFP channel (488 nm). Results were processed and visualized using FIJI. The lysates were washed off after each time point before imaging. At least 3,000 DNA molecules were imaged at each time point for each repeat.

### Optical trapping

#### Sample preparation

Cnn1 and Cnn1-3A purifications were performed similarly to the description above except for the removal of phosphatase inhibitors during rinses following the IP. Specifically, following the 3 hour IP, beads were washed three times with BH containing protease inhibitors and 2 mM dithiothreitol (DTT). Beads were then further washed twice with BH with protease inhibitors. The proteins were then eluted from the beads by gentle agitation in elution buffer (1 mg/ml 3xFlag peptide in BH) for 30 mins at room temperature, using 1/3 the bead volume. A small bead volume was used to enhance sample concentration for the trap.

For optical trapping, Dsn1 purifications were performed similar to Cnn1 purifications but without benomyl arrest or benzonase treatment to match prior purification protocols. Specifically, cultures were grown asynchronously in YPD to OD_600_ = 2-3, harvested, and washed once in cold water containing 0.2 mM PMSF (35 ml of water + PMSF for every 1 L culture harvested).

Washing, resuspension, and freezer milling were performed identically as for Cnn1 purifications, but no benzonase treatment was performed prior to ultracentrifugation. The IP was performed identically to Cnn1 except samples were eluted using 1/2 the bead volume.

#### Optical Trap Instrument

Optical trapping experiments were performed on a custom-built instrument based on designs described previously (*44*). Briefly, a trapping laser (Spectra-Physics; BL-106C), precision piezo-stage (Physik Instrumente; P-517.3CD), and position-sensitive photodetector (Pacific Silicon Sensor; DL100-7-PCBA2) were incorporated into an inverted light microscope (Nikon; Ti-U).

Prior to measurements, the position calibration of the trap was determined by raster scanning coverslip-bound beads and fitting photodetector response to a 5^th^-order polynomial. Trap stiffness was determined in 2 principal directions by averaging the stiffness determined from Stokes’ drag, equipartition, and power-spectrum fitting. Rupture force experiments were performed at similar stiffnesses for all bead diameters (*k*_x_∼0.16 pN/nm, *k*_y_∼0.1 pN/nm). A lower stiffness of *k*_x_∼0.03 pN/nm, *k*_y_∼0.2 pN/nm was used for constant force measurements near 2 pN. During measurement, bead-trap separation data and stage position data were sampled at 40 kHz, anti-alias filtered to 20 kHz (Krohn-Hite Model 3384), and decimated to 200 Hz before saving to disc. The force ramp measurement was controlled by custom LabView software (National Instruments), which updated the stage position at 50 Hz to achieve a force-ramp rate of 0.25 pN/s. Force clamp measurements used the same software with stage position used to maintain a constant set force near 2 pN.

#### Bead functionalization

Anti-His trapping beads were prepared from streptavidin-functionalized polystyrene beads. Dsn1-purification rupture forces and Cnn1purification force clamp experiments were performed with 350 nm average diameter beads (SVP-03-10, Spherotech). Additionally, Cnn1-purified kinetochores rupture force experiments in Fig. 7B, and binding experiments in fig. S2D were performed with 350 nm average diameter beads. For all other rupture force and bead binding experiments with Cnn1-purified kinetochores, 440 nm average diameter beads (SVP-05-10, Spherotech) were used. To prevent clumping and enhance homogeneous conjugation, beads were sonicated at 80% intensity in ice water for 5 minutes prior to functionalization (Biologics Inc.; Model 3000MP). Beads were functionalized for 1 hour at 4 °C by incubating beads at 0.1 % w/V with 50 µg/mL biotinylated anti-His antibody (R&D Systems; BAM050) in BRB80 (80mM K-PIPES, 1 mM MgCl_2_, 1 mM EGTA, pH = 6.9). Beads were rinsed 6 times by spinning down, removing supernatants, resuspending in 500 µL of BRB80 with 1 mM DTT (Fisher; BP172-5) and 8 mg/mL BSA (Millipore Sigma; A7906-10G), and sonicating for 1 minute at 58% intensity in ice water. Anti-His beads were stored rotating at 4 °C for 1-3 months. For experiments with unfunctionalized beads in Supplemental Figure 2A, streptavidin beads were prepared and rinsed identically to this protocol, without the addition of anti-His antibody.

#### Bead decorating

Kinetochores were linked to anti-His trapping beads by diluting purified kinetochores in assay buffer (BRB80, 2 mg/mL κ-Casein [Sigma-Aldrich C0406]) and then incubating with 0.01% w/V anti-His beads for 1 hour at 4 °C. Clumping was minimized by a 2-minute sonication of beads at 58% intensity in ice water prior to decoration and 30-s sonication at 25% intensity following the 1-hour incubation. Dsn1-purified kinetochores were diluted 20–25- fold to achieve ∼10-60% free-bead binding that has been previously shown yield consistent mean rupture forces (*42*). For rupture force experiments, Cnn1-purified kinetochores were diluted 2–10-fold resulting in varying extents of free bead binding (fig. S2B and S5). Because Cnn1-purified kinetochore rupture forces were similar across all bead dilutions, rupture force results were pooled from all bead dilutions. For constant force assays a dilution of 5-8-fold was used to achieve binding rates <25%, in an effort to achieve single particle binding. For bead binding assays in fig. S2A and S2D, Cnn1-purified kinetochore particles were diluted 2-fold to achieve high binding rates for comparison.

#### Dynamic microtubules

Optical trapping experiments were performed with commercial coverslips and slides cleaned for 5 minutes in a plasma cleaner (Harrick PDC-001). A small (∼3 mm wide) channel was formed between the slide and coverslip with double-sided tape (Scotch 662). Microtubule seeds were tethered to the coverslip surface by successive introduction of 10 µL 1 mg/mL biotinylated BSA (Pierce; 29130), 100 µL BRB80, 25 µL 1 mg/mL Avidin DN (Vector A-3100-1), 100 µL BRB80, and 100 µL GMPCPP-stabilized microtubule seeds. The remaining microtubule seeds were rinsed with 100 µL of growth buffer (BRB80, 1 mM GTP, 2 mg/mL K-Casein). Dynamic microtubules were grown from the coverslip by introduction of 25 µL of 15-20 µM bovine brain tubulin in trapping buffer with a glucose-oxidase oxygen scavenging system (BRB80, 1 mM GTP, 2 mg/mL K-Casein, 0.8 mM DTT, 20 mM glucose (Sigma-Aldrich; G8270), 125 U/mL glucose oxidase (Millipore Sigma; 345386), 420 U/mL catalase (Millipore Sigma; 209261), and 20 µg/mL biotinylated BSA). Trapping buffer included 20 µg/mL biotinylated BSA to block unconjugated streptavidin molecules on the beads and reduced sticking of beads to the coverslip.

#### Rupture force experiments

Kinetochore decorated beads were diluted ∼8.3-fold into trapping buffer and introduced to the channel. The slide was sealed and loaded onto the trap. Free beads were captured in the trap and positioned at microtubule tips to promote binding. Because the free bead binding rate tended to decrease as beads began to accumulate on microtubules, we only assessed free bead binding rates shortly following slide sealing. For fig. S2A and S2D, the binding rate for the first 25-30 beads were determined by the fraction of beads that could be bound to microtubules. For fig. S2B and S6, the binding rate for up to the first 20 free beads per slide was determined by the fraction of beads that could be bound to microtubules and could withstand a ∼1 pN preload once gently guided to the microtubule tip. The 1 pN preload helped screen for specific tip binding prior to initiating force ramps. Prebound beads that were found already bound to microtubules were used to measure rupture forces but not included in bead binding rates. Prebound beads were guided to the tip from their location on the microtubule and placed under a 1 pN preload. Beads that detached during preload were attempted a second time when possible before being discarded. Events that ended in detachment during preload were noted but not included in analysis.

Under preload, both free beads and prebound beads were assessed for stable tip attachment and displacement rates consistent with microtubule growth. Then a 0.25 pN/s force ramp was engaged to increase the applied force until the attachment ruptured, or the trace was interrupted by reaching the maximum force of the trap (∼25 pN for these experiments), microtubule detachment from the surface, or the trace being otherwise interrupted. Censored events accounted for <23% of analyzed traces in each experiment. Following rupture, the bead position at zero force was measured to provide an accurate zero for that trace. When necessary for censored events, another nearby bead would be used to measure the zero-force signal. Beads that ruptured from microtubules could often be rebound to another microtubule. A maximum of two events for each bead was included in analysis. Trapping slides were measured for up to 1.5 hours from sealing of the slide. Dsn1 particles rupture forces in Fig. 1 were performed from 2 distinct Cnn1 purifications, Cnn1 particles rupture forces in Fig. 1, Fig. 4, and Fig. 7A were performed from 4 distinct purification, Cnn1-3A particles rupture forces in Fig. 4 and Fig. 7A were performed from 3 purifications and mock treated Cnn1-3A particles rupture forces in Fig. 7A were performed from 2 purifications.

#### Stu2 Addback

For add back measurements, equal volume amounts of purified Stu2 or Stu2(Δ855-888) and purified Cnn1 or Cnn1-3A were incubated for 30 minutes rotating at 4 °C. These mixtures were then diluted with assay buffer and anti-His trapping beads to achieve similar final particle to bead dilutions as in normal rupture force experiments (typically 10%-45%) and incubated for 1 hr rotating at 4 °C. All other steps were identical to the purified Cnn1/Cnn1-3A rupture force experiments. For Stu2 add backs in Fig. 7A, Stu2-3Flag was purified from either asynchronously growing or benomyl arrested yeast cultures as described above. Because binding rates and rupture forces were similar for both forms of purified Stu2, results were pooled from both sources. Together these comprised three biological replicates of Stu2 add backs performed with multiple preparations of Cnn1 and Cnn1-3A purified kinetochore particles. For Stu2 add backs in Fig. 7B, Stu2 or Stu2(Δ855-888) were purified yeast cultures that were benomyl treated for 2.5 hours and treated with IAA for the final 60 minutes to deplete endogenous Stu2 as described above. Experiments for Fig. 7B comprise 2 replicates for both Stu2 or Stu2(Δ855-888) with a single preparation of Cnn1 and Cnn1-3A purified kinetochore particles. For mock experiments, Buffer H was used instead of purified Stu2.

#### Constant force experiments

Constant force trapping experiments were prepared similarly to rupture force experiments but performed using 350 nm beads and a lower trap stiffness as discussed above. As with rupture force measurements, both free beads and prebounds bead were used. Beads were guided to the microtubule tip and placed under a 1 pN preload and assessed for stable tip attachment. A 0.25 pN/s force ramp was engaged to increase the applied force to ∼2 pN. The stage-based feedback loop then held the bead force constant while tracking changes in the microtubule length corresponding to assembly and disassembly of microtubules. Events lasted until beads detached from the microtubule tip, or the event was otherwise interrupted. A zero-force measurement was made for each trace like in rupture force experiments. The length of events varied widely up to tens of minutes.

#### Trapping Data analysis

All optical trapping data was analyzed in Igor Pro (Wavemetrics) with custom written software (*44*). The zero-force stage position was determined from the trace following rupture, or from a nearby free bead if the interrogated bead could not be used. For constant force assays, the data was trimmed to the region held under constant force. For rupture force data, traces were discarded if the bead did not appear to rupture cleanly from the microtubule tip such as when bead position was translating rapidly, jumping or not stably associated with the microtubule within ∼1s of rupture, or if the rupture did not drop smoothly to zero force. Rupture points were determined individually by identifying the maximum force reached just prior to detachment of the bead from the microtubule. For censored data, the maximum force reached before the censoring event was used. Mean rupture forces were calculated from ruptured events and excluded any censored events. Error in the mean was calculated from the standard error of the mean. Statistical comparison of rupture force distributions was done with two-tailed Mann-Whitney U test in Igor Pro using the improved normal approximation. Survival curves were calculated from all events with the Kaplan-Meier estimator (*83*) in Igor Pro with 95% confidence intervals determined with log-log transformation. Survival curves were compared using the log-rank test. For statistical tests p<0.05 was considered significant.

The binding fraction (f) for each dilution was calculated by dividing the total number of beads that bound by the total number of free beads tested across one or more slides as described above. The uncertainty in the unbinding fraction (δf) was estimated from the Wald confidence interval for a binomial distribution using δf=√(f*(f-1))/N) where N is the total number of beads tested. For cases where no beads bound and the binding fraction was zero, we avoided the unrealistic δf = 0 by recalculating δf after adding a single bead binding event.

### Sequence alignment

The Cnn1 homolog sequences of other budding yeast species were obtained using NCBI BLAST and aligned using the MUSCLE algorithm in SnapGene.

### AlphaFold modeling

AlphaFold 3 multimer modeling (*84*) was used to predict the structure of the interaction between full-length Stu2 and 70 amino acids flanking the C-region of Cnn1, in a 2:1 stochiometric ratio. PyMOL (The PyMOL Molecular Graphics System, version 3.0, Schrodinger, LLC.) was used for visualization and manipulation of the structure.

## Acknowledgments

We thank members of the Biggins lab and Grant King for critical reading of the manuscript. We thank the proteomics and metabolomics core facility at Fred Hutch Cancer Center for the mass spectrometry sample processing and data analysis. We thank Trisha Davis, Arshad Desai, Elçin Űnal, Matthew Miller and Andrew Murray for providing reagents.

## Author contributions

Conceptualization: N.M., D.T.E., S.B.

Methodology: N.M., D.T.E., C.R.L., C.H., S.B.

Investigation: N.M., D.T.E., C.R.L., C.H., S.B.

Visualization: N.M., D.T.E., C.R.L., C.H., S.B.

Supervision: S.B.

Writing—original draft: N.M., D.T.E., S.B.

Writing—review & editing: N.M., D.T.E., C.L.A., S.B.

## Data, code, and materials availability

All data and code needed to evaluate and reproduce the results in the paper are present in the paper and/or the Supplementary Materials. Optical trapping results are provided in Tables S4-S6. Rupture force events and Kaplan Meier curves are provided in Table S4. Force clamp data and offsets used in creating plots for Fig. 1F are given in Table S5. Concentration-dependent optical trapping data including binding fractions, fits, and *F*_1/2_ data are given in Table S6. Raw traces are available. Mass spectrometry data presented in this study are available through Mass spectrometry interactive virtual environment (MassIVE, UCSD) https://massive.ucsd.edu/ProteoSAFe/private-dataset.jsp?task=8ce17ed601e343ae9edfcfbfa7f2fe92. The programs used for optical trapping analysis are available at https://github.com/casbury69/laser-trap-control-and-data-acquisition and https://github.com/casbury69/laser-trap-data-analysis.

