## Supplementary Material for "Kinetochore-microtubule attachments are strengthened by Cnn1 stabilization of Stu2"

Science Advances

AAAS

Nairita Maitra *et al.*

**This PDF file includes:**

Figs. S1 to S10
Tables S1 to S3

**Other Supplementary Materials for this manuscript include the following:**

Tables S4 to S6

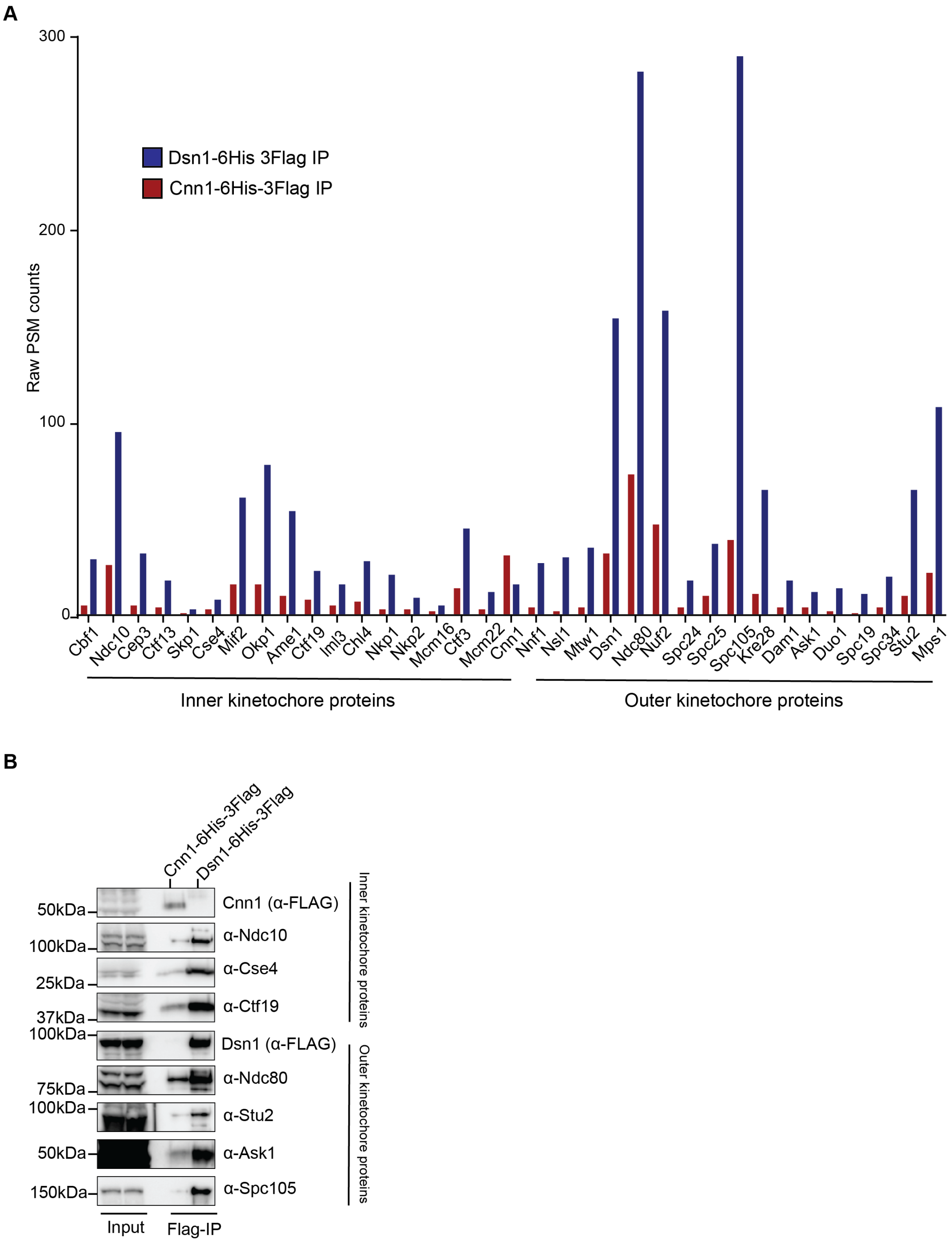


**Fig. S1. Kinetochore proteins that co-purify with Cnn1 and Dsn1.**

A. Mass spectrometry analysis of the kinetochore components from Cnn1 and Dsn1 purifications. Raw peptide spectral match (PSM) counts are presented in the y-axis.

B. Immunoblot analysis of kinetochore components in Cnn1-Flag or Dsn1-Flag purifications. Cells were arrested in mitosis with benomyl treatment and Cnn1-6His-3Flag (SBY24005) or Dsn1-6His-3Flag (SBY8253) was purified via anti-Flag immunoprecipitation.

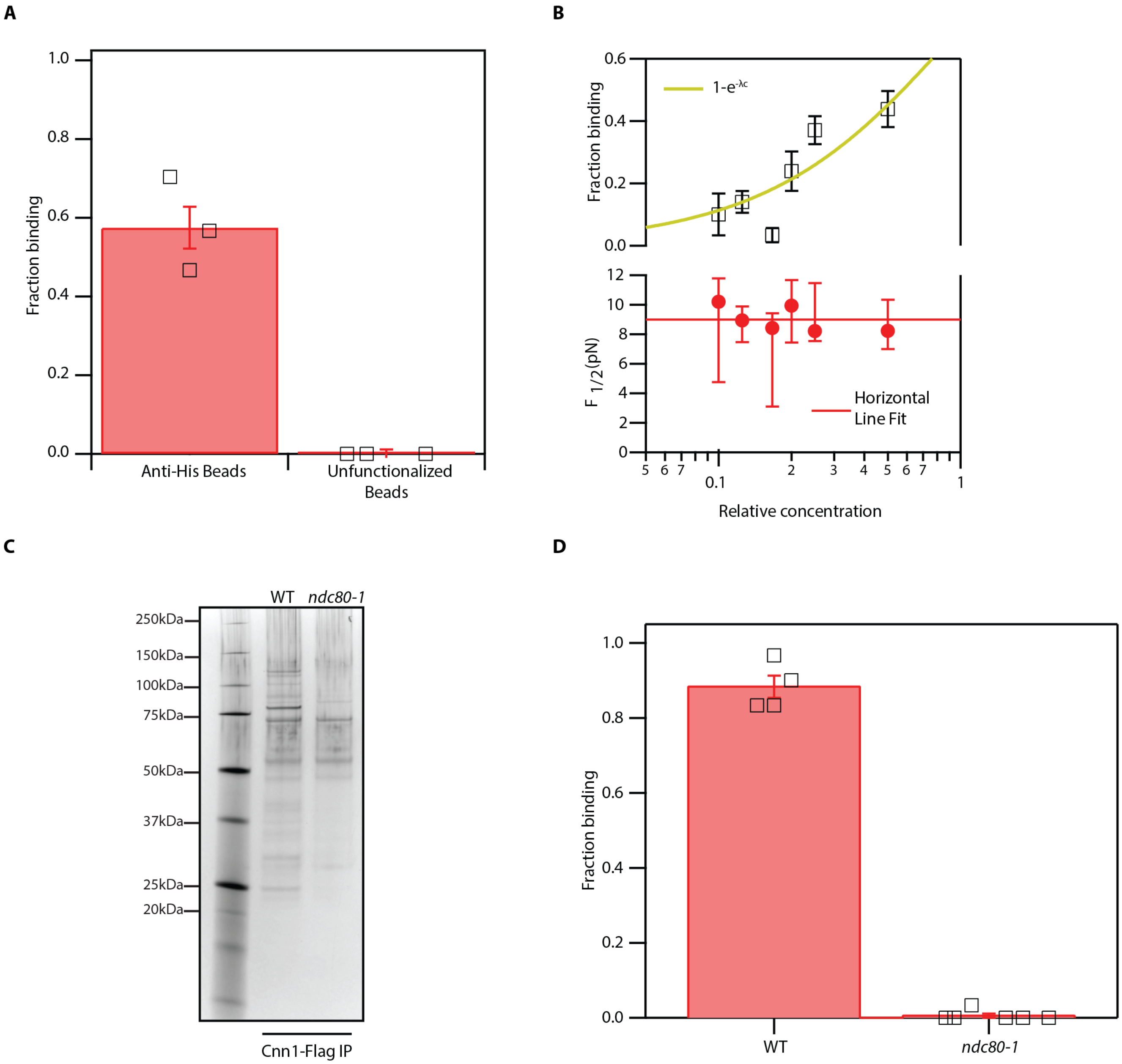


**Fig. S2. Rupture forces of Cnn1 particles are consistent with Ndc80-dependent single-particle microtubule binding.**

A. Fraction of free beads that bound microtubules in the optical trapping assay where Cnn1-6His-3Flag particles were incubated with either anti-His functionalized or unfunctionalized beads. Black squares show binding fractions measured on a single slide.

B. Top: Fraction of Cnn1-decorated beads that bound dynamic microtubules in optical trapping assay following incubation with varying concentrations of Cnn1 particles. The yellow line shows the binding likelihood assuming binding fractions are governed by the Poisson probability that a bead carries an active particle where c is the relative concentration and λ is a fitting parameter. The number of slides used for bead binding per dilution were: c = 0.5: 4 slides, c = 0.25: 6-slides, c = 0.2: 3 slides, c = 0.17: 3 slides; c = 0.125: 5 slides, c = 0.1: 1 slide. Bottom: The 50% survival force ($F_{1/2}$) for dilutions of Cnn1 purified kinetochores on beads. The error bar indicates 95% confidence intervals. The solid red line is a horizontal line fit to $F_{1/2}$. The number of events (n) per dilution was: c = 0.5: n = 117, c = 0.25: n = 96, c = 0.2: n = 58, c = 0.17: n = 14, c = 0.125: n = 82, c = 0.1: n = 11.

C. *CNN1-6His-3Flag* (SBY24005) and *CNN1-6His-3Flag ndc80-1* (SBY24285) cells were shifted to 37 °C for 3 hours in the presence of benomyl to arrest cells in mitosis. Cnn1 was immunoprecipitated and eluted with Flag peptide. The immunoprecipitations were analyzed by silver staining of SDS-PAGE.

D. Fraction of free beads that bound to microtubules in the optical trapping assay when Cnn1-6His-3Flag particles purified from *NDC80* or *ndc80-1* cells shifted to 37 °C were incubated with anti-His functionalized beads. Black squares show binding fractions measured on a single slide. Binding in cells with *NDC80* represents a single biological replicate, binding for *ndc80-1* cells is from 2 biological replicates.

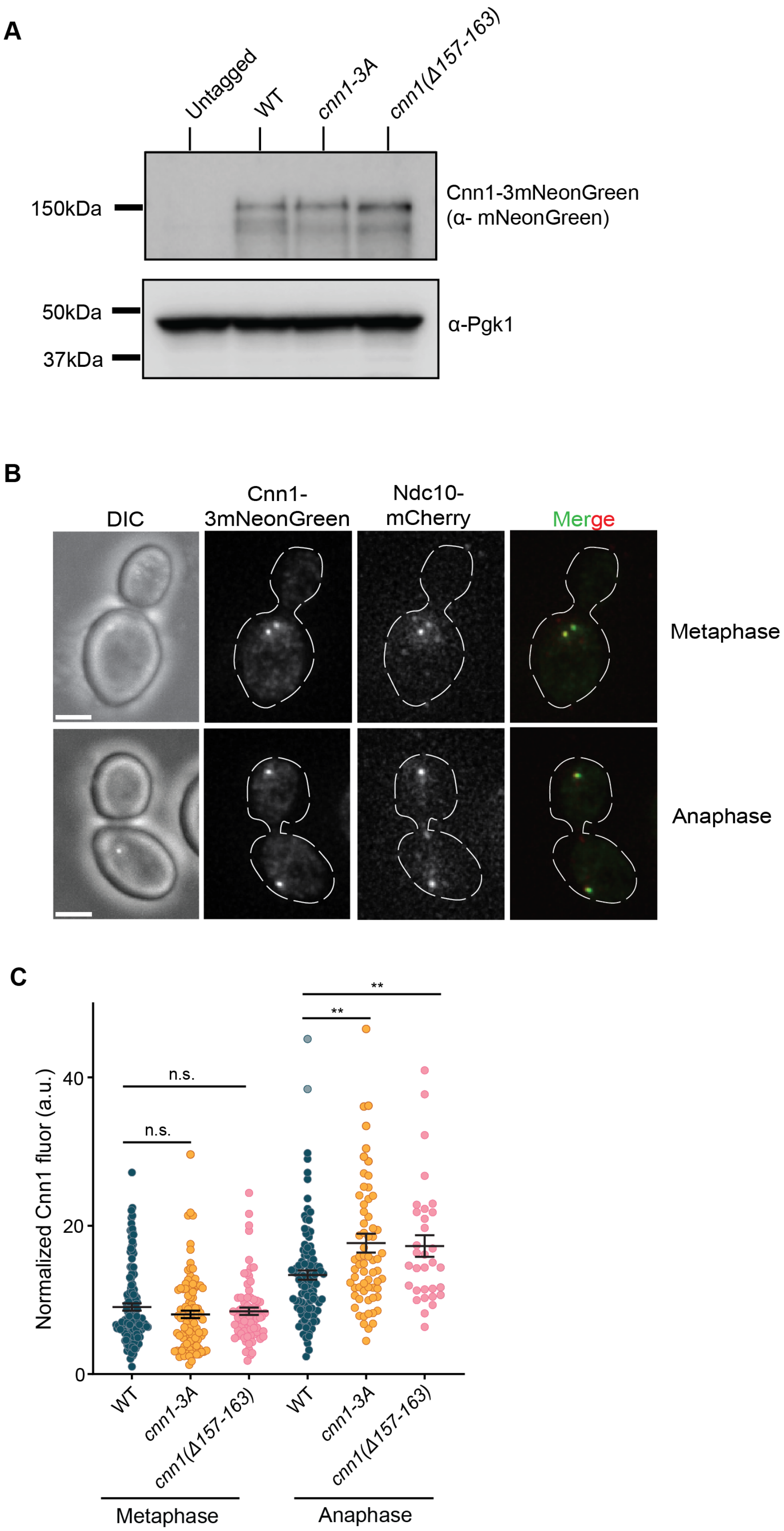

A
Untagged
WT
cnn1-3A
cnn1(Δ157-163)
150kDa
Cnn1-3mNeonGreen
(α- mNeonGreen)
50kDa
37kDa
α-Pgk1
B
DIC
Cnn1-
3mNeonGreen
Ndc10-
mCherry
Merge
Metaphase
Anaphase
C
Normalized Cnn1 fluor (a.u.)
0
20
40
n.s.
n.s.
**
**
WT
cnn1-3A
cnn1(Δ157-163)
WT
cnn1-3A
cnn1(Δ157-163)
Metaphase
Anaphase

**Fig. S3. Mutation of the Cnn1 C-region does not reduce Cnn1 protein levels.**

A. Cnn1-3mNeonGreen protein levels were assayed via immunoblot in *CNN1-3mNeonGreen* (SBY24029), *cnn1-3A-3mNeonGreen* (SBY24030), and *cnn1(Δ157-163)-3mNeonGreen* (SBY24031) cells from asynchronously growing cultures. Pgk1 was used as the loading control.
B. Representative images of Cnn1 kinetochore localization during metaphase and anaphase. Cells expressed endogenous Cnn1-3mNeonGreen and endogenous Ndc10-mCherry. Scale bar is 2 μm.
C. Quantification of Cnn1 levels at the kinetochore. *CNN1-3mNeonGreen* (SBY24029), *cnn1-3A-3mNeonGreen* (SBY24030), and *cnn1(Δ157-163)-3mNeonGreen* (SBY24031) cells were released from G1 and harvested at 75 mins and 100 mins to visualize metaphase and anaphase stages, respectively. Cnn1-3mNeonGreen fluor was quantified and normalized to the fiducial marker of Ndc10-mCherry. The p-values were calculated by Mann-Whitney test and for metaphase they are: WT vs *cnn1-3A* is 0.21; WT vs *cnn1(Δ157-163)* is 0.85; *cnn1-3A* vs *cnn1(Δ157-163)* is 0.31. For anaphase stage, the p-values are WT vs *cnn1-3A* is 0.002; WT vs *cnn1(Δ157-163)* is 0.007 and *cnn1-3A* vs *cnn1(Δ157-163)* is 0.942. The p-value between WT in metaphase vs WT in anaphase is < 0.0001. Error bars represent S.E.M. n.s. is non-significant. ** represents <0.01.

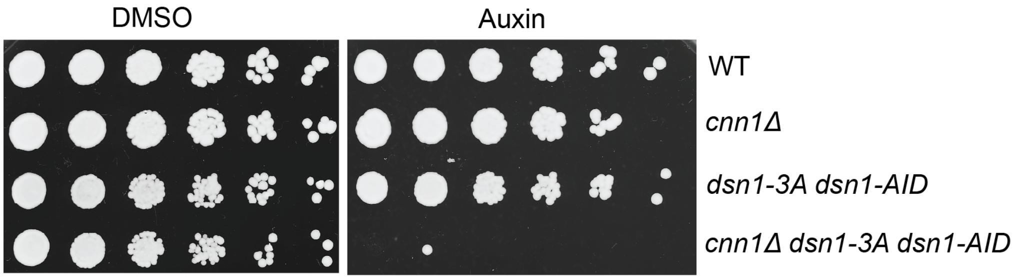


**Fig S4. *Cnn1Δ* cells are inviable when the Mis12c pathway is crippled.**

Viability of wild type (SBY3), *cnn1Δ* (SBY16332), *dsn1-3A dsn1-AID* (SBY19409), and *cnn1Δ dsn1-3A dsn1-AID* (SBY22387) cells was assayed via serial dilutions (5-fold dilutions) on YPD with either DMSO or 500 μM auxin.

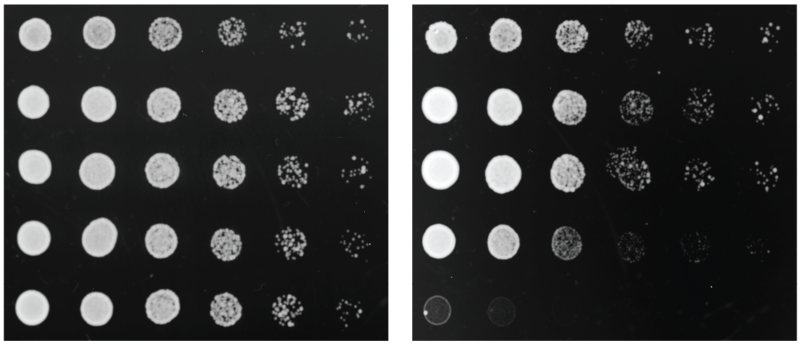


**Fig S5. *Cnn1-3A dsn1-3A* cells are inviable in absence of the *MAD3* spindle checkpoint gene.**

Viability of wild type (SBY3), *dsn1-3A dsn1-AID* (SBY22977), *cnn1-3A* (SBY24006) *cnn1-3A dsn1-3A dsn1-AID* (SBY22972) and *cnn1-3A dsn1-3A dsn1-AID mad3Δ* (SBY24389) cells was assayed via serial dilutions (5-fold dilutions) on YPD with either DMSO or 1mM auxin.

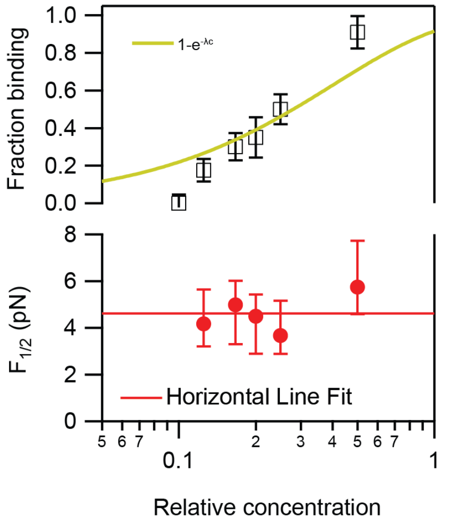


**Fig S6. Rupture forces of Cnn1-3A particles are consistent with single-particle microtubule binding.**

Top: Fraction of Cnn1-3A particle-decorated beads that bound dynamic microtubules in optical trapping assay following incubation with varying concentrations of Cnn1-3A purifications. The yellow line shows the binding likelihood assuming binding fractions are governed by the Poisson probability that a bead carries an active particle where c is the relative concentration and λ is a fitting parameter. The number of slides used for bead binding at each relative concentration are: c = 0.5: 1 slide, c = 0.25: 2-slides, c = 0.2: 1 slide, c = 0.17: 2 slides, c = 0.125: 2 slides, c = 0.1: 1 slide. Bottom: The 50% survival force ($F_{1/2}$) for dilutions of Cnn1-3A purified kinetochores on beads. The error bar indicates 95% confidence intervals. The solid red line is a horizontal line fit to $F_{1/2}$. The number of events (n) per dilution were: c = 0.5: n = 30, c = 0.25: n = 52, c = 0.2: n = 22, c = 0.17: n = 34, c = 0.125: n = 37, c = 0.1: n = 0.

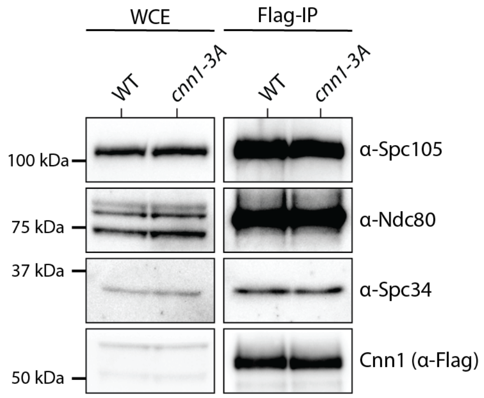


**Fig S7. Outer kinetochore composition analysis of Cnn1 and Cnn1-3A purifications.**

Cnn1-3Flag (SBY24005) or Cnn1-3A-3Flag (SBY24006) were purified from cells arrested in mitosis by benomyl treatment. Outer kinetochore protein levels were analyzed by immunoblot

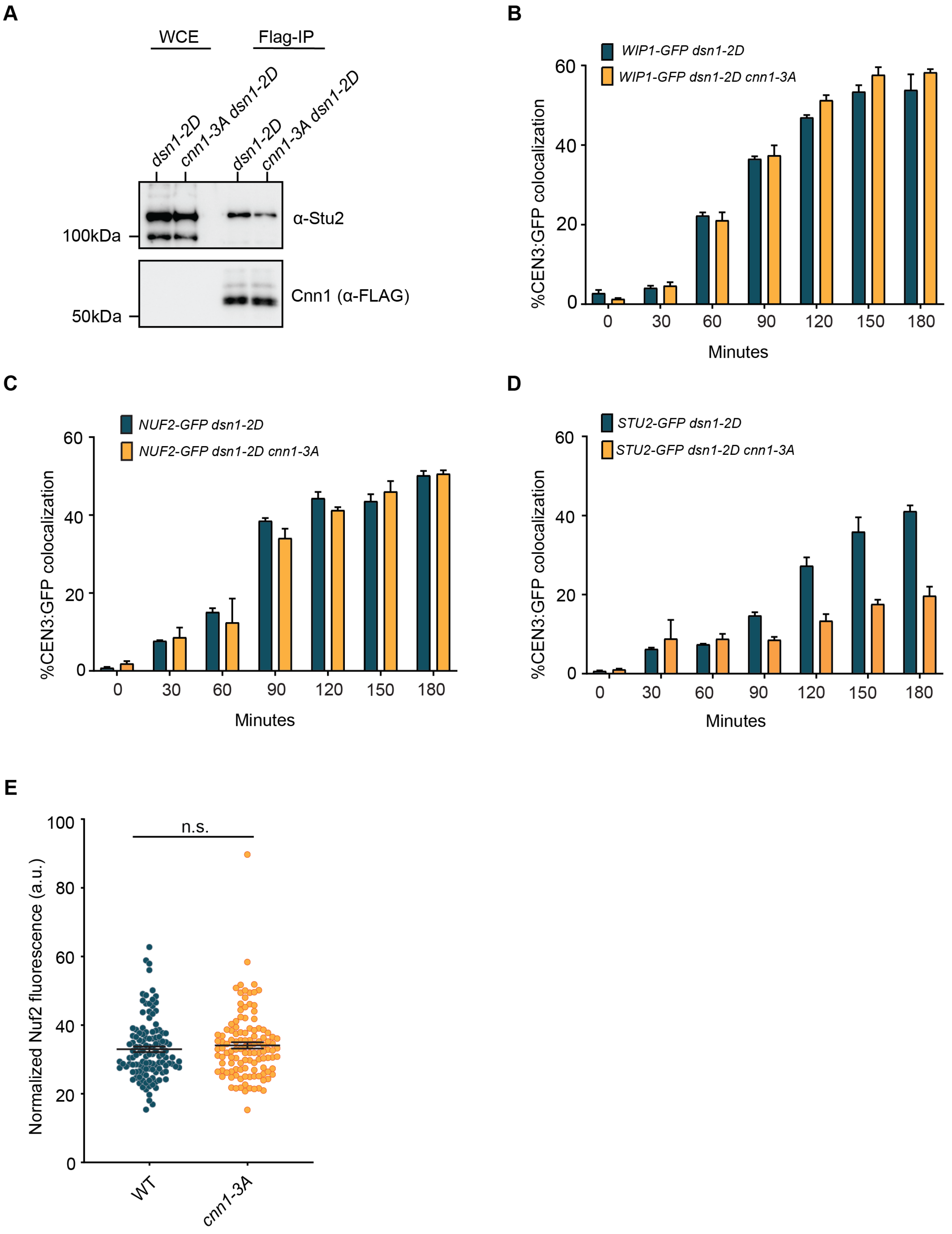
A
WCE
Flag-IP
dsn1-2D
cnn1-3A dsn1-2D
dsn1-2D
cnn1-3A dsn1-2D
α-Stu2
100kDa
Cnn1 (α-FLAG)
50kDa
B
WIP1-GFP dsn1-2D
WIP1-GFP dsn1-2D cnn1-3A
%CEN3:GFP colocalization
Minutes
C
NUF2-GFP dsn1-2D
NUF2-GFP dsn1-2D cnn1-3A
%CEN3:GFP colocalization
Minutes
D
STU2-GFP dsn1-2D
STU2-GFP dsn1-2D cnn1-3A
%CEN3:GFP colocalization
Minutes
E
n.s.
Normalized Nuf2 fluorescence (a.u.)
WT
cnn1-3A

**Fig S8.** ***Cnn1-3A*** **cells exhibit reduced Stu2 levels at the kinetochore.**

A. Cnn1-Flag was immunoprecipitated from benomyl arrested *dsn1-2D* (SBY23855) and *dsn1-2D cnn1-3A* (SBY23856) cells. Stu2 levels were analyzed via immunoblot.
B. Percentage of Atto-647 *CEN3* DNA colocalized with Wip1-GFP measured by single molecule TIRF assay using *WIP1-GFP dsn1-2D* (SBY22205) and *WIP1-GFP dsn1-2D cnn1-3A* (SBY24956) lysates. 3000 DNA molecules were imaged at each time point for each strain and biological replicate (n = 3). The time axis represents the incubation time of the DNA with the lysate before it was washed out and TIRFM was performed. The p-value calculated from T-test over three experiments at 180 mins is 0.044.
C. Percentage of Atto-647 tagged *CEN3* DNA colocalized with Nuf2-GFP measured by a single molecule TIRF assay using *NUF2-GFP dsn1-2D* (SBY23258) and *NUF2-GFP dsn1-2D cnn1-3A* (SBY24955) cells. 3000 DNA molecules were imaged at each time point for each strain and biological replicate (n = 3). The time axis represents the incubation time of the DNA with the lysate before it was washed out and TIRFM was performed. The p-value calculated from T-test over three experiments at 180 mins is 0.8715.
D. Percentage of Atto-647 *CEN3* DNA colocalized with Stu2-GFP measured by single molecule TIRF assay using *STU2-GFP dsn1-2D* (SBY22133) and *STU2-GFP dsn1-2D cnn1-3A* (SBY24954) cells. 3000 DNA molecules were imaged at each time point for each strain and biological replicate (n = 3). The time axis represents the incubation time of the DNA with the lysate before it was washed out and TIRFM was performed. The p-value calculated from T-test over three experiments at 180 mins is <0.0001.
E. Quantification of Nuf2 level at kinetochores *in vivo*. Wild type cells (SBY24827) and *cnn1-3A* cells (SBY24826) containing Nuf2-GFP and Ame1-mKATE were released from G1 and harvested 75 mins after release to obtain metaphase cells. The fluor intensity of Nuf2-GFP was quantified and normalized to the Ame1-mKate intensity. The p-value (0.323) is calculated from the Mann-Whitney test across 2 experiments (n > 100 cells for each strain). The error bar represents S.E.M. n.s. represents non-significant.

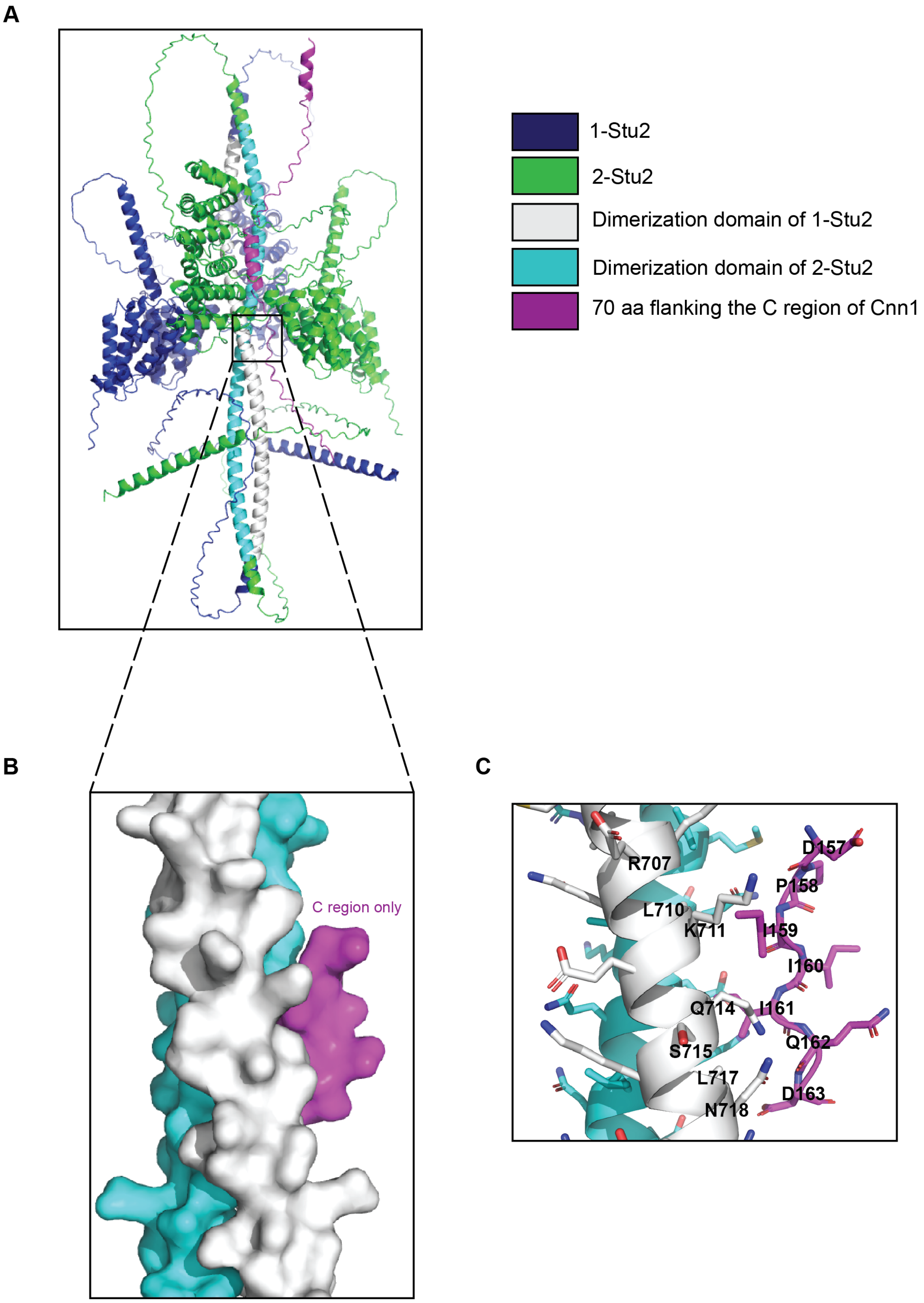

A
1-Stu2
2-Stu2
Dimerization domain of 1-Stu2
Dimerization domain of 2-Stu2
70 aa flanking the C region of Cnn1
B
C region only
C
D157
R707
P158
L710
K711
I159
I160
Q714
I161
S715
Q162
L717
D163
N718

**Fig S9. AlphaFold3 modeling of possible interaction between Stu2 and the C region of Cnn1**.

A. AlphaFold 3 multimer model of 2 Stu2 molecules with 70 amino acids (106 - 178 amino acids) of Cnn1 flanking the C region. The Stu2 and the C region of Cnn1 are color-coded, as shown in the right panel. Blue represents 1 molecule of Stu2, and Green represents the second molecule of Stu2. The dimerization domains of the first and second Stu2 molecules are color-coded in cyan and grey, respectively. The Stu2 dimerization domains have high confidence (pIDDT>90), while the C region has low confidence (pIDDT<50). The overall interaction model has iPTM =0.27 and pTM = 0.32.

B. Zoomed in view of (A) to highlight the Stu2 dimerization interface and the C region of Cnn1. The protein surface of the interaction region is shown.

C. The residues of Stu2 at the dimerization interface potentially interacting with the C region of Cnn1 are displayed.

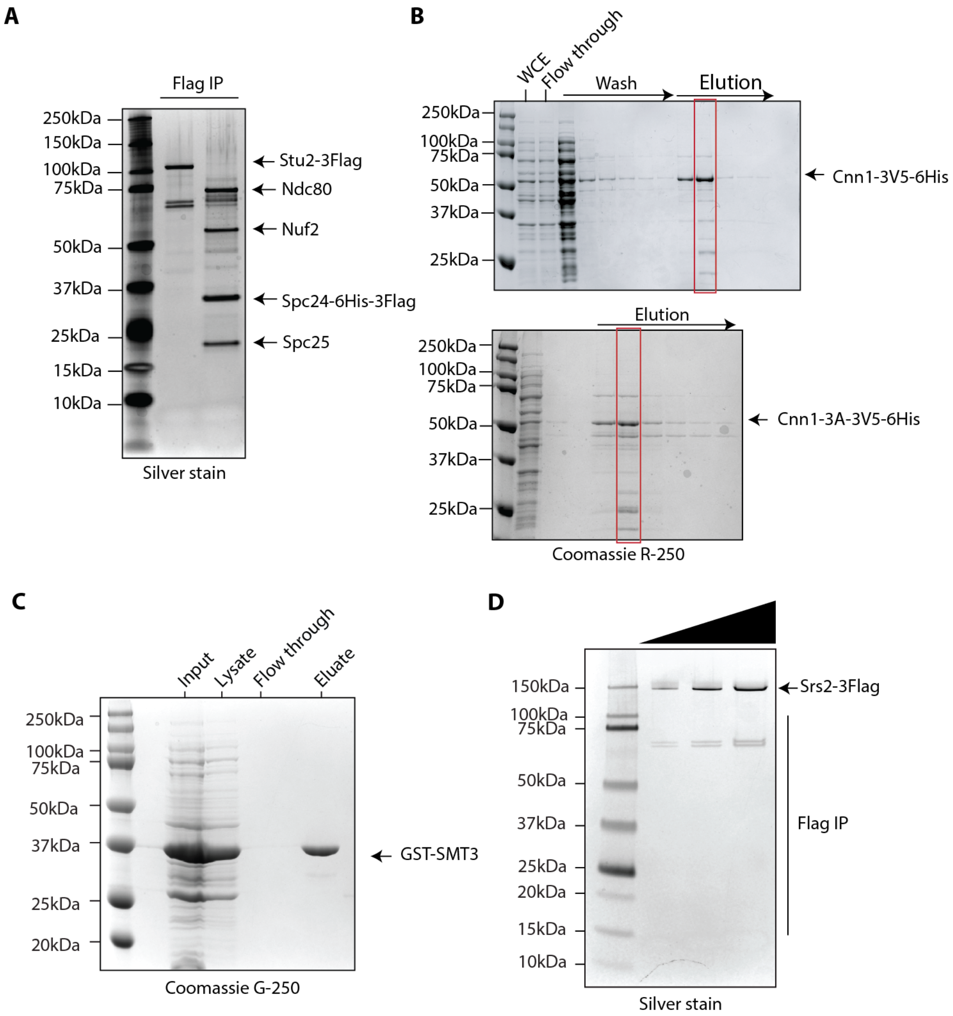


**Fig S10. Analysis of purification of Stu2, Ndc80c, Srs2 and recombinant Cnn1 and Cnn1-3A**

A. High salt purifications of Stu2-Flag (SBY12275) (left lane) and Ndc80c via Spc24-Flag (SBY14022) (right lane) from budding yeast cells arrested in mitosis by benomyl treatment were analyzed by silver staining of SDS-PAGE.
B. Coomassie staining of SDS-PAGE gels analyzing recombinant wild type Cnn1 (top panel) and Cnn1-3A (bottom panel) proteins expressed in *E. coli* and purified using immobilized metal ion affinity chromatography. Eluate 2, shown in the red box, was used for both Cnn1 and Cnn1-3A binding assays shown in panel C.
C. Coomassie staining of SDS-PAGE gels analyzing recombinant GST-Smt3 and GST proteins expressed in *E. coli* (GST-Smt3: pSB3783 and GST: pSB3778) were purified by affinity purification using glutathione sepharose resins.
D. High salt purification of Srs2-Flag (SBY25556) from budding yeast cells was analyzed by silver staining of SDS-PAGE.

### Supplementary Tables

### Table S1: List of strains used in this study

| Strain | Genotype |
|---|---|
| SBY3 (W303) | *MATa ura3-1 leu2-3,112 his3-11 trp1-1 can1-100 ade2-1 bar1-1* |
| SBY8253 | *MATa DSN1-6His-3FLag:URA3* |
| SBY8315 | *MATα NDC10-mCherry:HPH* |
| SBY11586 | *MATa CNN1-3Flag:KANMX6* |
| SBY12275 | *MATa STU2-3FLag:KANMX6* |
| SBY14022 | *MATa SPC24-6His-3Flag:URA3 SPC105-3HA-IAA7:KANMX6 trp1-1::pGPD1-OsTIR1:TRP1* |
| SBY16332 | *MATa cnn1Δ::HIS3* |
| SBY19409 | *MATa his3-11::dsn1-3A:HIS3 leu2-3,112::pGPD1-OsTIR1:LEU2 DSN1-3HA-IAA7:KANMx6* |
| SBY21592 | *MATa leu2-3,112::GFP-TUB1:LEU2 MTW1-mCherry:HPH* |
| SBY21682 | *MATa CNN1-3Flag:KANMX6 STU2-3V5:HIS3MX6* |
| SBY21805 | *MATa cnn1-3A-3Flag:KANMX6 STU2-3V5:HIS3MX6* |
| SBY21908 | *MATa leu2-3,112::GFP-TUB1:LEU2 MTW1-mCherry:HPH cnn1-3A* |
| SBY22078 | *MATa cnn1-3A-3FLag:KANMX6* |
| SBY22133 | *MATα STU2-GFP:KANMX6 dsn1-2D-3Flag:URA3* |
| SBY22174 | *MATa STU2-GFP:KANMX6* |
| SBY22205 | *MATα WIP1-GFP:KANMX6 dsn1-2D-3Flag:URA3* |
| SBY22242 | *MATα AME1-mKATE2:HIS3MX6* |
| SBY22387 | *MATa his3-11::dsn1-3A:HIS3 leu2::pGPD1-OsTIR1:LEU2 DSN1-3HA-IAA7:KANMX6 cnn1Δ::HIS3* |
| SBY22830 | *MATa his3-11::dsn1-3A:HIS3 trp1-1:pGPD1-OsTIR1:TRP1 leu2-3,112::GFP-TUB1:LEU2 DSN1-3HA-IAA7:KANMX6 MTW1-mCherry:HPH* |
| SBY22831 | *MATa his3-11::dsn1-3A:HIS3 trp1-1:pGPD1-OsTIR1:TRP1 leu2-3,112::GFP-TUB1:LEU2 DSN1-3HA-IAA7:KANMX6 MTW1-mCherry:HPH cnn1-3A* |
| SBY22972 | *MATa his3-11::dsn1-3A:HIS3 leu2-3,112::pGPD1-OsTIR1:LEU2 DSN1-3HA-IAA7:KANMX6 cnn1-3A-6His-3FLag:URA3* |
| SBY22977 | *MATa his3-11::dsn1-3A:HIS3 leu2-3,112::pGPD1-OsTIR1:LEU2 DSN1-3HA-IAA7:KANMX6 CNN1-6His-3FLag:URA3* |
| SBY23258 | *MATa NUF2-mGFP:KANMX6 dsn1-2D-3Flag:URA3* |
| SBY24005 | *MATa CNN1-6His-3Flag:URA3* |
| SBY24006 | *MATa cnn1-3A-6His-3Flag:URA3* |
| SBY24029 | *MATa NDC10-mCherry:HPH CNN1-3mNeonGreen:HIS3* |
| SBY24030 | *MATa NDC10-mCherry:HPH cnn1-3A-3mNeonGreen:HIS3* |
| SBY24031 | *MATa NDC10-mCherry:HPH cnn1(Δ157-163)-3mNeonGreen:HIS3* |

| | |
|---|---|
| SBY24336 | *MATa ura3-1::pCyc1-GFP12-LacI12:URA3 his3-11::dsn1-3A:HIS3 chr8::CEN256LacO:TRP1 leu2-3,112::pGPD1-OsTIR1:LEU2 DSN1-3HA-IAA7:KANMX6 mad3Δ::NATMX6* |
| SBY24387 | *MATa PDS1-13Myc:LEU2 CNN1-6His-3Flag:URA3* |
| SBY24389 | *MATa his3-11::dsn1-3A:HIS3 leu2-3,112::pGPD1-OsTIR1:LEU2 DSN1-3HA-IAA7:KANMX6 cnn1-3A-6His-3Flag:URA3 mad3Δ::NATMX6* |
| SBY24391 | *MATa mad3Δ::NATMX6* |
| SBY24542 | *MATa ura3-1::pCyc1-GFP12-LacI12:URA3 chr8::CEN256LacO:TRP1 Cnn1-6His-3FLag:URA3 mad3Δ::NATMX6* |
| SBY24544 | *MATa ura3-1::pCyc1-GFP12-LacI12:URA3 chr8::CEN256LacO:TRP1 cnn1-3A-6His-3FLag:URA3 mad3Δ::NATMX6* |
| SBY24549 | *MATa PDS1-13Myc:LEU2 his3-11::dsn1-3A:HIS3 leu2-3,112::pGPD1-OsTIR1:LEU2 DSN1-3HA-IAA7:KANMX6 cnn1-3A-6His-3Flag:URA3 mad3Δ::NATMX6* |
| SBY24552 | *MATa cnn1(Δ157-163)-6His-3Flag:URA3* |
| SBY24565 | *MATa ura3-1::pCyc1-GFP12-LacI12:URA3 his3-11::dsn1-3A:HIS3 chr8::CEN256LacO:TRP1 leu2-3,112::pGPD1-OsTIR1:LEU2 DSN1-3HA-IAA7:KANMX6 cnn1-3A-6His-3Flag:URA3 mad3Δ::NATMX6* |
| SBY24652 | *MATa his3-11::dsn1-3A:HIS3 leu2-3,112::pGPD1-OsTIR1:LEU2 DSN1-3HA-IAA7:KANMX6 cnn1(Δ157-163)-6His-3Flag:URA3* |
| SBY24738 | *MATa PDS1-13Myc:LEU2 CNN1-6His-3Flag:URA3 mad3Δ::NATMX6* |
| SBY24741 | *MATa STU2-GFP:KANMX6 cnn1-3A-6His-3Flag:URA3 AME1-mKATE2:HisMX6* |
| SBY24782 | *MATa STU2-GFP:KANMX6 CNN1-6His-3Flag:URA3 AME1-mKATE2:HIS3MX6* |
| SBY24789 | *MATa PDS1-13Myc:LEU2 his3-11::dsn1-3A:HIS3 leu2-3,112::pGPD1-OsTIR1:LEU2 DSN1-3HA-IAA7:KANMX6 cnn1-3A-6His-3Flag:URA3 mad3Δ::NATMX6* |
| SBY24826 | *MATa NUF2-mGFP:KANMX6 cnn1-3A-6HIS-3FLAG:URA AME1-mKate2:HIS3MX6* |
| SBY24827 | *MATa NUF2-mGFP:KANMX6 CNN1-6HIS-3FLAG:URA AME1-mKate2:HISMX6* |
| SBY24828 | *MATa TOR1-1 fprΔ::NATMX6 STU2-FRB:HIS3 NUF2-2FKBP12:TRP1 CNN1-6His-3Flag:URA3* |
| SBY24829 | *MATa TOR1-1 fprΔ::NATMX6 STU2-FRB:HIS3 CNN1-6His-3Flag:URA3* |
| SBY24830 | *MATa TOR1-1 fprΔ::NATMX6 STU2-FRB:HIS3 NUF2-2FKBP12:TRP1 cnn1-3A-6His-3Flag:URA3* |
| SBY24831 | *MATa TOR1-1 fprΔ::NATMX6 STU2-FRB:HIS3 cnn1-3A-6His-3Flag:URA3* |
| SBY24833 | *MATa TOR1-1 fprΔ::NATMX6 STU2-FRB:HIS3 NUF2-2FKBP12:TRP1 his3-11::dsn1-3A:HIS3 leu2-3,112::pGPD1-OsTIR1:LEU2 DSN1-3HA-IAA7:KANMX6 CNN1-6His-3Flag:URA3* |

| | |
|---|---|
| SBY24835 | *MATa TOR1-1 fprΔ::NATMX6 STU2-FRB:HIS3 NUF2-2FKBP12:TRP1 his3-11::dsn1-3A:HIS3 leu2-3,112::pGPD1-OsTIR1:LEU2 DSN1-3HA-IAA7:KANMX6 cnn1-3A6His-3Flag:URA3* |
| SBY24900 | *MATa TOR1-1 fprΔ::NATMX6 STU2-FRB:HIS3 his3-11::dsn1-3A:HIS3 leu2-3,112::pGPD1-OsTIR1:LEU2 DSN1-3HA-IAA7:KANMX6 CNN1-6His-3Flag:URA3* |
| SBY24902 | *MATa TOR1-1 fprΔ::NATMX6 STU2-FRB:HIS3 his3-11::dsn1-3A:HIS3 leu2-3,112::pGPD1-OsTIR1:LEU2 DSN1-3HA-IAA7:KANMX6 cnn1-3A-6His-3Flag:URA3* |
| SBY24954 | *MATα STU2-GFP:KANMX6 dsn1-2D-3Flag:URA3 cnn1-3A* |
| SBY24955 | *MATα NUF2-mGFP:KANMX6 dsn1-2D-3Flag:URA3 cnn1-3A* |
| SBY24956 | *MATα WIP1-GFP:KANMX6 dsn1-2D-3Flag:URA3 cnn1-3A* |
| SBY25530 | *MATa cnn1-(Δ65-79)-6HIS-3FLAG:URA3 Stu2-3V5:HisMX* |
| SBY25556 | *MATα SRS2-3FLAG:HPHMX6* |
| SBY25566 | *MATa his3-11:dsn1-3A:HIS3 pGPD1-OsTIR1:LEU DSN1-3HA-IAA7:KANMx6 CNN1-HIS-FLAG:URA3 TOR1-1 fpr1Δ::NATMx STU2-FRB:HIS NUF2-2FKB12:TRP MTW1-mCherry:HPH* |
| SBY25569 | *MATa his3-11:Dsn1-3A:HIS3 pGPD1-OsTIR1:LEU DSN1-3HA-IAA7:KANMx6 CNN1-3A-6HIS-3FLAG:URA TOR1-1 fpr1Δ::NATMx STU2-FRB:HIS NUF2-2FKB12:TRP MTW1-mCherry:HPH* |
| SBY25588 | *Mata leu2-3,112::pSTU2-Stu2(Δ855-888)-3Flag:LEU2 his3::pGPD1-OsTIR1:HIS3 STU2-3HA-IAA7:KanMX* |
| SBY25592 | *Mata leu2-3,112::pSTU2-STU2-3Flag:LEU2 his3::pGPD1-OsTIR1:HIS3 STU2-3HA-IAA7:KanMX* |

**Table S2: List of plasmids used in this study**

| Plasmid # | Purpose |
|---|---|
| pSB812 | For Gibson assembly generation of pSB3779 and pSB3780. 3Flag-KANmX replaced 3V5 in pSB2358 and pSB2232. |
| pSB1108 | To integrate *dsn1-3A* at the ectopic locus of *HIS3*. |
| pSB1590 | To epitope tag *CNN1* with 6His-3Flag at its C-terminus. |
| pSB2232 | Used as template for *STU2 f*or Gibson assembly generation of pSB3780 . |
| pSB2358 | Used as template for *stu2(Δ855-888) f*or Gibson assembly generation of pSB3779. |
| pSB3405 | URA3-CEN-sgRNA-Cas9 plasmid targeting ATACAAGATGTTCCCACGAC on *CNN1* to create *cnn1-3A* mutation by CRISPR. |
| pSB3524 | Used as a template to create plasmids pSB3667 and pSB3740. |
| pSB3575 | To epitope tag *CNN1* with 3mNeonGreen at the C-terminus. |
| pSB3667 | To induce expression of recombinant Cnn1-3V5-6His in bacteria by IPTG. |
| pSB3740 | To induce expression of recombinant Cnn1-3A-3V5-6His in bacteria by IPTG. |
| pSB3741 | URA-CEN-sgRNA-Cas9 plasmid targeting ATAATAATACAAGATGTTCC on *CNN1* to create *cnn1(Δ157-163)* by CRISPR. |
| pSB3776 | GST-SMT3 bacterial expression (SUMO mature form SMT3gg) |
| pSB3779 | To integrate *stu2(Δ855-888)-3FLAG::LEU2* at the *leu2* locus |
| pSB3780 | To integrate *STU2-3FLAG::LEU2* at the *leu2* locus |

**Table S3: List of primers used in this study (5' to 3')**

| Oligos | Purpose | Sequence |
|---|---|---|
| SB660 | Reverse primer to sequence *HIS3* locus to verify *dsn1-3A* integration | GTTTAAGGCTAGTGAGTGCG |
| SB4007 | Forward primer to amplify *CNN1* (starting at 184bp) to confirm *cnn1-3A* or *cnn1(Δ157-163)* mutations | GATCCGAACGAAGTACGAAG |
| SB4126 | Reverse primer (~300bp downstream of STOP codon of *CNN1*) to check the 6His-3Flag integration at C-terminus of *CNN1* | ACACTCGTGGCAATGACTCAG |
| SB4265 | Reverse primer to amplify *CNN1* (250bp upstream of the stop codon of *CNN1*) to confirm *cnn1-3A* or *cnn1(Δ157-163)* mutations | AGATCCATTATTTGCAGCGG |
| SB4819 | Forward primer for confirming integration of *STU2* and *stu2(Δ855-888)* at *LEU2* locus | gcagattccctttatggattcc |
| SB4820 | Forward primer for confirming integration of *STU2* and *stu2(Δ855-888)* at *LEU2* locus | ggtagatttagtactgaagaggaggtcg |
| SB5178 | Forward primer (~300bp upstream of STOP codon of *CNN1*) to check the 6His-3Flag integration at C-terminus of *CNN1* | CCGTTTAAATGTCCCTGCGG |
| SB7632 | Forward primer to clone *CNN1* sgRNA into Cas9 plasmid to create *cnn1-3A* | ggctgggcaacaccttcgggtggcgaatggATACAAGATGTTCCCACGAC |
| SB7633 | Reverse primer to clone *CNN1* sgRNA into Cas9 plasmid to create *cnn1-3A* | attttaacttgctatttctagctctaaaacGTCGTGGGAACATCTTGTAT |
| SB7634 | Repair template to create *cnn1-3A* mutation by CRISPR | TGGCAATGAAACGCAACCTGATTATACTTCCTTATCGCAAACAGTATTTGCAAAATTGCAGGAAAGAGACAAAGGCTTGAAAAGTAGGAAGATTGAC |

| | | |
|---|---|---|
| | | CCGgcagcagcaCAAGATGTTCCCACGACAGGACACGAGGATGAATTGACAGTGCACTCTCCAGATAAAGCTAACAGTATCTCTATGGAAGTTTTGCGCACGT |
| SB8093 | Forward primer to sequence *HIS3* locus to verify *dsn1-3A* integration | GATGTTCCCTCCACCAAAG |
| SB8711 | Forward primer to clone *CNN1* sgRNA into Cas9 plasmid to create *cnn1(Δ157-163)* | ggctgggcaacaccttcgggtggcgaatggATAATAATACAAGATGTTCC |
| SB8712 | Reverse primer to clone *CNN1* sgRNA into Cas9 plasmid to create *cnn1(Δ157-163)* | attttaacttgctatttctagctctaaaacGGAACATCTTGTATTATTAT |
| SB8713 | Repair template to create *cnn1(Δ157-163)* mutation by CRISPR | CAACATTGGCAATGAAACGCAACCTGATTATACTTCCTTATCGCAAACAGTATTTGCAAAATTGCAGGAAAGAGACAAAGGCTTGAAAAGTAGGAAGATTGTTCCCACGACAGGACACGAGGATGAATTGACAGTGCACTCTCCAGATAAAGCTAACAGTATCTCTATGGAAGTTTTGCGCACGTCGCCAAGCATTGGAA |
| SB8800 | Reverse primer to epitope tag *CNN1* with 3-mNeonGreen at its C-terminus | TTTTACTCTTTTCTGTACATAAATTGCCACTATTTAATTATTTTCTCTACGGTATCTTTTggcgttagtatcgaatcgacagc |
| SB8801 | Forward primer to epitope tag *CNN1* with 3-mNeonGreen at its C-terminus | CAACATTGGCAATGAAACGCAACCTGATTATACTTCCTTATCGCAAACAGTATTTGCAAAATTGCAGGAAAGAGACAAAGGCTTGAAAAGTAGGAAGATTGTTCCCACGACAGGACACGAGGATGAATTGACAGTGCACTCTCCAGATAAAGCTAACAGT |

| | | |
|---|---|---|
| | | ATCTCTATGGAAGTTTTGCGCACGTCGCCAAGCATTGGAA |
| SB8883 | Forward primer to amplify vector pSB3524 | GAATTGAAAGTTATTTGTTCcggatccccgggttaattaacatct |
| SB8884 | Reverse primer to amplify vector pSB3524 | GCCTTCCTGGGAGTGCTCATTGTATATCTCCTTCTTAAAGTTAAACaaaattatttctagaggg |
| SB8885 | Forward primer to amplify *CNN1* from gDNA | CTTTAAGAAGGAGATATACAATGAGCACTCCCAGGAAGG |
| SB8886 | Reverse primer to amplify *CNN1* from gDNA | ttaattaacccggggatccgGAACAAATAACTTTCAATTCTTGATTGAAGTTCTAAGGG |
| SB9069 | Forward primer to epitope tag *CNN1* with 6His-3Flag:URA | GAGCTCGCGCACGAATTGCTTCCCTTAGAACTTCAATCAAGAATTGAAAGTTATTTGTTCgAATTCaagcttgggttaattaac |
| SB9070 | Reverse primer to epitope tag *CNN1* with 6His-3Flag:URA | TTTTACTCTTTTCTGTACATAAATTGCCACTATTTAATTATTTTCTCTACGGTATCTTTTgattcggtaatctccgaacag |
| SB9561 | Forward primer for Gibson assembly of pSB3779 from pSB812 | TAGTTTTGCCAGGGAACAAAAGCTGGAG |
| SB9562 | Reverse primer for Gibson assembly of pSB3779 from pSB812 | TAGTGGATCCTAGCCGCGTTCTAACGAC |
| SB9563 | Forward primer for Gibson assembly of pSB3779 from pSB2358 | AACGCGGCTAGGATCCACTAGTTCTAGAC |
| SB9564 | Reverse primer for Gibson assembly of pSB3779 from pSB2358 | TTTGTTCCCTGGCAAAACTATCGTTGAAATC |
| SB9565 | Forward primer for Gibson assembly of pSB3780 from pSB812 | AACCAGGACGAGGGAACAAAAGCTGGAG |
| SB9566 | Reverse primer for Gibson assembly of pSB3780 from pSB812 | TAGTGGATCCTAGCCGCGTTCTAACGAC |

| | | |
|---|---|---|
| SB9567 | Forward primer for Gibson assembly of pSB3780 from pSB2232 | AACGCGGCTAGGATCCACTAGTTCTAGACAC |
| SB9568 | Reverse primer for Gibson assembly of pSB3780 from pSB2232 | TTTGTTCCCTCGTCCTGGTTGTCCCTTC |

**Other supplementary materials:**

Table S4: Rupture force events and Kaplan Meier curves – provided in excel file

Table S5: Force clamp data and offsets – provided in an excel file

Table S6: Binding rate data – provided in an excel file
